# Testing for Shared Molecular-Evolutionary Signatures of Diurnality and Nocturnality in Mammalian Circadian Clock Genes

**DOI:** 10.64898/2026.09.17.752369

**Authors:** Alper Kaan Selçukoğlu, Ali Koray Koç

## Abstract

Following an ancestral nocturnal phase, diurnality evolved independently in multiple mammalian lineages after the Cretaceous–Palaeogene boundary. Whether these recurrent temporal-niche transitions drove convergent molecular evolution in the circadian oscillator remains largely untested. We analyzed 18 core clock genes and direct regulators across 60 diurnal and nocturnal mammals, identifying 10 independent gains of diurnality and 5 reversals to nocturnality on a fixed supertree topology. Using six complementary evolutionary frameworks (spanning substitution counts, profile shifts, selective regimes, and evolutionary rates) we tested for shared molecular adaptations. To ensure that our negative findings reflected biological reality rather than methodological insensitivity, we planted 90 synthetic convergent residues into the empirical alignments. While this benchmark demonstrated high power to detect broadly shared convergence (e.g., PCOC recovered 35 of 36 planted sites across ten lineages without false positives), empirical alignments showed no credible signal. Same-residue convergence occurred at expected background levels (0.310 observed vs. 0.337 null), and no gene displayed activity-dependent shifts in selective pressure or evolutionary rate. These results indicate that repeated transitions in mammalian activity patterns were not driven by a common set of detectable amino-acid changes in the core circadian machinery.

## Introduction

Evidence supports a nocturnal origin of mammals and a prolonged nocturnal phase during the Mesozoic, with diurnality emerging mainly after the Cretaceous–Palaeogene extinction (Gerkema et al., 2013; Maor et al., 2017). Diurnality subsequently evolved independently in several mammalian lineages. These repeated transitions provide an opportunity to test whether similar molecular changes accompanied independent shifts towards the same activity pattern. However, similar phenotypes can evolve through different molecular routes, so sequence convergence is a hypothesis to be tested rather than a necessary consequence of phenotypic convergence (Natarajan et al., 2016).

The mammalian circadian clock generates approximately 24-hour rhythms through interacting transcription–translation feedback loops (Partch et al., 2014; Takahashi, 2017). CLOCK or NPAS2 pairs with BMAL1, encoded by ARNTL, to activate transcription at E-box elements. PER1 and PER2 act with CRY1 and CRY2 to repress this activity, whereas PER3 contributes to circadian timing in a more tissue-dependent manner (Pendergast et al., 2012). An additional feedback loop regulates ARNTL transcription through ROR response elements: REV-ERBα/β, encoded by NR1D1 and NR1D2, repress transcription, whereas RORα/β/γ, encoded by RORA, RORB and RORC, activate it. Protein modification and degradation also help determine circadian period. The casein kinases CK1δ and CK1ε, encoded by CSNK1D and CSNK1E, phosphorylate PER proteins and regulate their stability and localisation, while FBXL3 promotes ubiquitin-dependent degradation of CRY proteins. BHLHE40 and BHLHE41, encoding DEC1 and DEC2, provide additional repression of CLOCK–BMAL1-dependent transcription (Honma et al., 2002). We analysed these 18 genes as a targeted panel of core clock components and direct regulators to test whether repeated transitions between diurnal and nocturnal activity are associated with sequence changes in the circadian machinery.

Associations between circadian-gene variation and activity timing have been documented most extensively within species. In humans, for example, a coding variable-number tandem repeat in PER3 has been associated with extreme diurnal preference and delayed sleep phase, while large genome-wide association studies have identified numerous loci associated with chronotype (Archer et al., 2003; Jones et al., 2019). Comparative evidence across mammals remains more limited and has generally focused on particular ecological transitions, individual lineages or restricted groups of genes. Examples include analyses of PER and CRY genes in subterranean rodents (Sun et al., 2018) and a genome-wide study of the origin of diurnality in the striped mouse Rhabdomys pumilio (Richardson et al., 2023). The expectation that temporal-niche evolution can leave a coding-sequence signature also has precedent in the mammalian visual system. Comparative analyses of opsins have identified gene losses consistent with prolonged nocturnal ancestry, as well as activity-associated differences at functionally important spectral-tuning sites (Borges et al., 2018). These findings concern the sensory machinery that detects light, however, rather than the circadian oscillator itself. To our knowledge, the protein-coding sequences of the core mammalian clock have not previously been tested in a taxon sample designed around multiple independent transitions to diurnality and reversals to nocturnality.

Because identical amino acids frequently arise through mutation biases and substitution rate heterogeneity, identifying repeated substitutions alone is insufficient to establish adaptive convergence. The same amino acid can arise independently because of mutation and fixation biases, variation in substitution rate or the limited number of possible amino-acid states. Repeated substitutions must therefore be compared with the amount of convergence expected in the absence of a phenotype-associated effect (Rey et al., 2019). The importance of this null expectation was illustrated by reanalyses of echolocating mammals: an apparent genome-wide excess of convergent substitutions between bats and dolphins did not exceed the background observed in appropriately matched comparisons (Thomas and Hahn, 2015; Zou and Zhang, 2015). A second difficulty is phylogenetic non-independence. Diurnal and nocturnal species are not distributed randomly across the mammalian tree, so conventional parametric probabilities can be miscalibrated unless the null preserves the phylogenetic structure of the phenotype (Saputra et al., 2021). Finally, different methods test different forms of molecular similarity: repeated changes to the same residue, shifts in amino-acid preference, changes in site-specific selective pressure and changes in evolutionary rate are related but not interchangeable signals. We therefore used six complementary analyses and required support from more than one method for a site-level signal to be considered high confidence.

To meaningfully interpret a null finding, the analytical pipeline must demonstrate adequate statistical power to detect the target signal if present. Statistical power in convergence analysis is not determined by the number of phenotypic transitions alone; it also depends on the topology and branch lengths of the tree, alignment properties and the strength and distribution of the molecular signal. Power must therefore be evaluated for the dataset and convergence scenario under study rather than inferred from sample size alone (Rey et al., 2018). Simulations under a detection method’s own evolutionary model are useful for characterising method-specific behaviour, but they cannot provide an end-to-end check or reveal an incorrectly constructed convergence scenario, mislabelled transition branches or an error introduced earlier in the processing pipeline. We therefore complemented method-specific calibration by planting known convergent residues into the empirical alignments and rerunning the site-level analyses without changing the trees, scenario files, scripts or decision thresholds. Recovery of planted sites and detection of unmodified sites were recorded separately. This design allows a negative result to be interpreted in terms of the classes and strengths of convergence that the pipeline demonstrably recovers.

Here, we test whether the molecular evolution of 18 circadian genes tracks mammalian diel activity beyond the similarity expected from shared ancestry. We analyse 60 mammal species selected for clear diurnal or nocturnal classification and to represent repeated evolutionary transitions between these states. We ask whether species sharing an activity pattern have more similar sequences than their phylogenetic relationships predict, whether independently shifted lineages experienced comparable changes in evolutionary rate or selective regime, and whether independent transitions involved convergent amino-acid substitutions at homologous sites. These questions are addressed using six complementary approaches across 10 independent gains of diurnality and 5 reversals to nocturnality identified under the primary equal-rates ancestral-state reconstruction. The two directions of change are analysed as separate convergent classes because they lead towards opposite phenotypic states.

## Materials and Methods

### Taxon sampling and phenotype data

Sixty mammalian species were sampled across the placental orders together with a marsupial outgroup. Diel activity was coded as a binary character from published compilations (Maor et al. 2017; Bennie et al. 2014), giving 30 diurnal and 30 nocturnal species. No sampled species carried an intermediate crepuscular or cathemeral score, so no intermediate states were collapsed. Taxa were chosen to maximise the number of independent transitions between activity states while retaining phylogenetic spread. This yields a diel sample that is deliberately balanced and therefore does not reflect the nocturnal predominance of mammals as a whole; the consequences are addressed in the ancestral state reconstruction below. Species were selected on the joint availability of behavioural evidence and of genomic resources suitable for recovering orthologous sequences of the target genes, so the sample is not a random draw from mammalian temporal diversity and is biased towards lineages with better genomic coverage.

Every activity assignment was re-evaluated against the primary literature. Each species was coded by the phase containing the majority of its activity. Where a crepuscular component was documented it was assigned to the adjacent light or dark phase, according to whether activity was weighted towards dawn or towards dusk and night. For species with seasonal, thermal, disturbance-related or population-level plasticity, the dominant modal pattern was used and the qualification recorded. Thirty-two assignments are unambiguous and 28 are dominant-phase and qualified; confidence is high for 42 species, medium for 15 and low for 3 (Table S1). Four domesticated taxa were retained and flagged separately, because their observed activity timing may partly reflect husbandry.

### Target genes and protein alignments

Protein sequences were collected for 18 core circadian genes spanning the functional modules of the transcription-translation feedback loop: the positive arm (CLOCK, NPAS2, ARNTL), the negative arm (PER1, PER2, PER3, CRY1, CRY2), the auxiliary loop (NR1D1, NR1D2, RORA, RORB, RORC), post-translational regulators (CSNK1D, CSNK1E, FBXL3) and the output repressors (BHLHE40, BHLHE41). Protein sequences were recovered and passed through orthology assessment and protein-level quality control before any coding sequence was reconstructed. Three additional genes considered during panel construction were not retained. ARNTL2 was initially included, but reciprocal sequence searches and orthology checks did not consistently distinguish ARNTL2 from ARNTL across the sampled taxa, with ARNTL repeatedly recovered in place of ARNTL2. ARNTL2 was therefore removed before alignment construction and hypothesis testing. TIMELESS and FBXL21 were not retained under the final targeted-panel definition. Mammalian TIMELESS is more closely related to the ancestral *timeout/tim2* lineage than to *Drosophila timeless* and has major functions in replication-fork stability and genome maintenance (Gotter et al., 2007; Cai et al., 2022). FBXL21, although relevant to circadian regulation, has compartment-dependent effects on CRY stability that can oppose or complement those of FBXL3; the targeted panel represented this CRY-turnover system with FBXL3 (Hirano et al., 2013; Yoo et al., 2013). The resulting 18-gene set should therefore be regarded as a targeted panel of core clock components and selected direct regulators rather than a complete inventory of genes associated with circadian regulation. Taxon occupancy is incomplete for some genes; per-gene alignments contain between 40 and 60 species (Table 1).

**Table 1.**
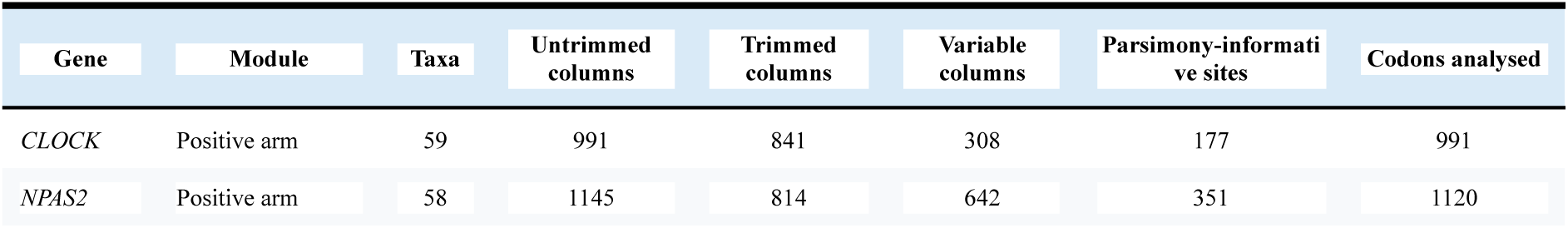
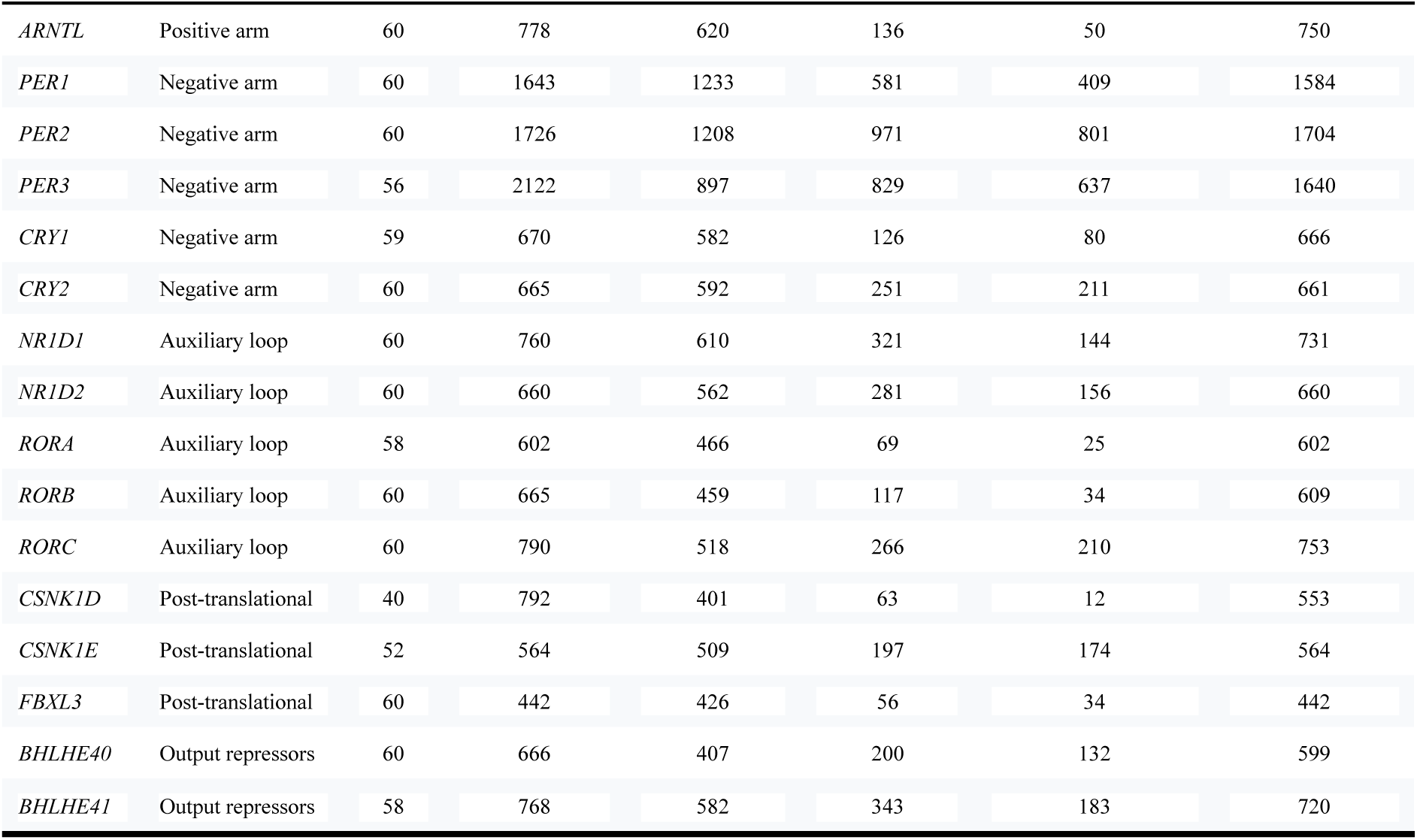
Gene set, taxon occupancy and alignment dimensions. Trimmed columns are those retained by trimAl-automated1. A column is variable when it holds more than one amino acid, and parsimony-informative when at least two amino acids each occur in at least two sequences. The parsimony-informative columns are exactly the sites TDG09 could test; it returns NA at every other column. Codons analysed are those remaining after removal of all-gap columns.

| Gene | Module | Taxa | Untrimmed columns | Trimmed columns | Variable columns | Parsimony-informative sites | Codons analysed |
| --- | --- | --- | --- | --- | --- | --- | --- |
| CLOCK | Positive arm | 59 | 991 | 841 | 308 | 177 | 991 |
| NPAS2 | Positive arm | 58 | 1145 | 814 | 642 | 351 | 1120 |
| *ARNTL* | Positive arm | 60 | 778 | 620 | 136 | 50 | 750 |
| *PER1* | Negative arm | 60 | 1643 | 1233 | 581 | 409 | 1584 |
| *PER2* | Negative arm | 60 | 1726 | 1208 | 971 | 801 | 1704 |
| *PER3* | Negative arm | 56 | 2122 | 897 | 829 | 637 | 1640 |
| *CRY1* | Negative arm | 59 | 670 | 582 | 126 | 80 | 666 |
| *CRY2* | Negative arm | 60 | 665 | 592 | 251 | 211 | 661 |
| *NR1D1* | Auxiliary loop | 60 | 760 | 610 | 321 | 144 | 731 |
| *NR1D2* | Auxiliary loop | 60 | 660 | 562 | 281 | 156 | 660 |
| *RORA* | Auxiliary loop | 58 | 602 | 466 | 69 | 25 | 602 |
| *RORB* | Auxiliary loop | 60 | 665 | 459 | 117 | 34 | 609 |
| *RORC* | Auxiliary loop | 60 | 790 | 518 | 266 | 210 | 753 |
| *CSNK1D* | Post-translational | 40 | 792 | 401 | 63 | 12 | 553 |
| *CSNK1E* | Post-translational | 52 | 564 | 509 | 197 | 174 | 564 |
| *FBXL3* | Post-translational | 60 | 442 | 426 | 56 | 34 | 442 |
| *BHLHE40* | Output repressors | 60 | 666 | 407 | 200 | 132 | 599 |
| *BHLHE41* | Output repressors | 58 | 768 | 582 | 343 | 183 | 720 |

Alignment headers were verified against species tree tip labels before any downstream analysis. Alignments were trimmed with trimAl v1.5.rev1 (Capella-Gutierrez et al. 2009) under the-automated1 heuristic, retaining for each gene a map from every trimmed column to its index in the untrimmed alignment. That map, combined with the *Homo sapiens* alignment row, converts trimmed column indices to human residue numbering for reporting. The trimmed dataset comprises 11,727 columns, of which 5,970 are constant, 5,757 variable and 3,820 parsimony-informative (Table 1). The parsimony-informative columns are exactly the sites TDG09 was able to test.

### Coding-sequence recovery

Synonymous and nonsynonymous substitutions cannot be distinguished from protein alignments, so the codon-level analyses reported here, Contrast-FEL and RELAX in HyPhy (Wertheim et al. 2015; Kosakovsky Pond et al. 2021), require coding-sequence alignments, as do the wider family of codon methods that this dataset is therefore able to support. We recovered the coding sequence corresponding to each protein retained in the protein alignments.

Recovery followed two principles. First, each CDS had to correspond to the exact protein record used in the protein alignment rather than to the same gene and species, because retrieving a different isoform can produce a codon alignment inconsistent with the protein dataset. Second, codon alignments were generated from the existing protein alignments with pal2nal (Suyama et al. 2006) rather than by aligning nucleotide sequences independently, which preserves the reading frame and gives protein-level and codon-level analyses a common coordinate system.

The complete design comprises 1,080 species by gene combinations. After protein retrieval and protein-level quality control, 1,040 sequences remained: 15 combinations could not be filled and 25 sequences were excluded during protein-level filtering. These 1,040 proteins defined the target set for CDS recovery, of which 593 were assigned to accession-based retrieval and 447 to genome-based reconstruction; the species by gene provenance map is given in Table S2.

For proteins with annotated NCBI accessions, the CDS was recovered from the accession of the exact aligned protein rather than through a gene-name search. GenPept records were retrieved through NCBI Entrez using Biopython v1.84 (Cock et al. 2009) and the /coded_by qualifier was parsed for the corresponding nucleotide accession and CDS coordinates, after which the specified interval was retrieved directly. For records with missing or complex /coded_by expressions, including joined or complemented intervals that could not be parsed reliably, a fallback used elink to connect the protein record to nuccore; candidate coding sequences were retrieved in fasta_cds_na format and matched back to the original protein accession through the protein_id field. This route is called accession-based rather than RefSeq-based because it also covers annotated GenBank protein accessions.

Twenty-six species were represented by unannotated GenBank assemblies and lacked suitable annotated CDS records; their protein sequences had been predicted with miniprot v0.18 (Li 2023) from multi-species query pools, with the selected hit, query protein and target contig recorded for each species by gene combination. For CDS reconstruction, miniprot was rerun in --gff mode using the query protein recorded at the protein-recovery step, which preserves provenance and yields GFF3 records with explicit CDS coordinates. Where several hits were returned, the hit on the originally selected contig was preferred. CDS features were ordered in transcript direction, reverse-complemented for minus-strand predictions and concatenated, and the leading phase of the first CDS feature was applied before translation. Because rerunning miniprot does not guarantee that the original prediction is reproduced, reconstructed sequences were subjected to the same verification as accession-derived sequences.

One systematic taxon-label mismatch was corrected at this stage. NCBI records for aardvark use the trinomial Orycteropus afer afer whereas the alignment tips use Orycteropus afer, which initially prevented accession resolution for all 17 aardvark records. After normalisation, every sequence in the target set was assigned to one of the two retrieval routes.

### CDS verification and codon alignments

Accession-derived and genome-reconstructed sequences were verified identically. Each CDS was translated under the standard genetic code (NCBI translation table 1), all 18 genes being nuclear, and the translation was compared with the ungapped protein sequence used in the corresponding protein alignment. A CDS was retained only if its translation matched the target protein in length, contained no internal stop codons and agreed at every residue, with an X in the target protein allowed to match any translated amino acid. Sequences failing these criteria were excluded and assigned an explicit failure reason. Only verified sequences were passed to pal2nal. For each gene the protein alignment was restricted to species with a verified CDS and ordered identically to the nucleotide input; codon alignments were exported in FASTA format for HyPhy and in PAML format for codon-model software that requires it.

Of the 1,040 target sequences, 823 (79.1%) passed verification and entered the codon alignments. On the accession-based route 9 of 593 sequences were lost (1.5%): eight failed verification on internal stop codons and one, RORB in Rhynchocyon petersi, could not be retrieved at all because the elink fallback failed repeatedly for that record. Genome-based reconstruction failed for 208 of 447 sequences (46.5%), so genome-predicted sequences account for 208 of the 217 losses at this stage (95.9%). The most frequent failure categories were internal stop codons, disagreement between CDS and protein length, and CDS lengths not divisible by three. These patterns are compatible with errors in gene prediction, genome assembly or CDS reconstruction, but they do not identify the cause of any individual failure: verification excluded sequences inconsistent with the protein dataset without assigning each discrepancy a single cause.

### Codon-level coverage

Codon-level coverage ranges from 32 species for CSNK1D to 58 for FBXL3, with a median of 45. To test whether attrition differed between activity classes, the diurnal and nocturnal composition of each codon alignment was compared with the original 30:30 design. Balance ratios, the size of the smaller class divided by that of the larger, range from 0.77 to 1.00 and mostly exceed 0.90. ARNTL, BHLHE40, BHLHE41 and CRY1 retain equal numbers of diurnal and nocturnal species and NR1D2 retains a 28:28 split. Five genes fall below 40 species at the codon level: CSNK1D (32), PER3 (36), BHLHE40 (38), CSNK1E (39) and RORB (39). Numerical balance between activity classes does not exclude phylogenetically structured missingness, so coverage was considered together with the number of diel-state changes each gene retains.

One species, *Nanger dama* (diurnal, Bovidae), is absent from all 18 codon alignments and was excluded from codon-level analyses. Ten further species occur in fewer than half of the codon alignments: its sister tip in the reference topology, *Aepyceros melampus,* in 4 of 18; *Caracal caracal, Daubentonia madagascariensis, Nasua narica, Paguma larvata, Pipistrellus pipistrellus* and *Tapirus terrestris* in 7 of 18; and *Choloepus hoffmanni, Mirza zaza* and *Speothos venaticus* in 8 of 18. With *Nanger dama* these eleven low-coverage species comprise seven nocturnal and four diurnal taxa, so attrition among the repeatedly missing taxa is skewed towards the nocturnal class even though individual alignments remain close to balanced. All eleven were represented only by the genome-reconstruction route. Their uneven phylogenetic distribution was handled by evaluating retained diel-state changes separately for each gene.

CSNK1D has the lowest codon-level coverage, but most of that reduction happened before CDS verification: 20 of its 60 protein sequences had already been excluded at protein-level quality control, including predictions of 737 aa (*Acomys cahirinus*) and 744 aa (*Hylobates pileatus*) against a reference length near 415 aa. The remaining 40 sequences entered verification as 32 accession-based and 8 miniprot-derived records. All 32 accession-based sequences passed; all 8 genome-derived sequences failed on low identity, internal stops, lengths not divisible by three or disagreement with the target protein length. The final alignment therefore contains 32 species. No CDS inconsistent with its protein sequence entered any codon alignment.

Protein-level analyses retain more species and were used wherever the broadest taxon coverage was wanted; codon-level analyses were restricted to sequences passing CDS to protein verification. The two datasets are treated as complementary because they retain different species and support different classes of analysis.

### Retention of diel-state transitions

Species count alone does not measure the number of independent phenotypic transitions available to a convergence analysis, because several closely related tips may represent one transition. To describe the transition replication retained after sequence filtering, the reference topology (Upham et al. 2019) was pruned separately for each gene and Fitch parsimony (Fitch, 1971) was used to count the minimum number of diurnal to nocturnal changes required on each pruned tree. The complete 60-species tree has a minimum parsimony score of 14 state changes. Gene-specific scores range from 9 for CSNK1D to 14 for FBXL3 and NR1D2, with ARNTL at 11 and PER3 at 10. Taxon counts and parsimony scores were kept as separate descriptors of the available data.

Fitch parsimony gives the minimum number of changes compatible with the observed tip states. It does not incorporate branch lengths, unequal transition rates or repeated changes along one branch, so the score of a pruned tree describes minimum transition replication and is not a formal estimate of statistical power. It also does not establish that the same historical transitions survived pruning, since alternative equally parsimonious reconstructions may place changes on different branches.

### Phylogenetic framework

The topology was pruned from the mammalian supertree of Upham et al. (2019) and held fixed for every analysis, so that transition branches occupy identical nodes across all 18 genes. Branch lengths were estimated in IQ-TREE v3.1.1 (Wong et al. 2026) under this fixed topology (-te), which optimises lengths without topology search. Where a gene lacked taxa present in the species tree, the tree was pruned to that gene’s taxa before estimation.

Two branch length sets were produced. Analyses treating genes independently (PCOC, TDG09, the selection tests) used per-gene best-fit models selected by ModelFinder (Kalyaanamoorthy et al. 2017). RERconverge, which compares rates across genes, used a single uniform model (Q.MAMMAL+F+R6) selected by running ModelFinder once on the concatenation of all 18 trimmed alignments; differing rate-heterogeneity models across genes would otherwise introduce systematic differences in branch length shape unrelated to biology. Mean branch length spans a 37-fold range across genes (0.0035 substitutions per site in ARNTL to 0.130 in CSNK1E), which is why detection power was calibrated per gene rather than once for the dataset.

Because IQ-TREE returns unrooted trees while the species tree is rooted on the marsupial outgroup, and the two rootings induce different clade decompositions near the root, every per-gene tree was re-rooted on the outgroup before scenario construction.

Unconstrained per-gene maximum likelihood trees and a partitioned supermatrix tree were estimated separately and used only for orthology quality control, concordance factor estimation and topology confirmation. They were not used as an analysis backbone.

### Ancestral state reconstruction and definition of convergent events

Ancestral diel states were reconstructed with corHMM v2.8 (Boyko and Beaulieu, 2021) using marginal reconstruction, one rate category and a free root. Phylogenetic signal in the trait was assessed with Pagel’s λ in phytools v2.5.2 (Revell, 2024).

Two models were fitted and compared by Akaike information criterion (AIC) as returned by the corHMM fit, with one free rate parameter under equal rates and two under all rates different. The equal-rates (ER) model had the lower AIC (ER 77.82 against ARD 79.27, delta AIC = 1.45) and was designated the primary analysis on that basis before any convergence detection was run. That margin is narrow enough that the two reconstructions are close to equally supported, which is why the all-rates-different (ARD) model was retained as a sensitivity analysis on the reconstruction. Fixing the primary reconstruction in advance is what allows a site detected only under ARD to be set aside without a post hoc choice. Because gains of diurnality and reversals to nocturnality move toward opposite phenotypes, they were analysed as separate convergent classes throughout; PCOC fits a single convergent amino acid profile per run, so merging them would be self-cancelling.

Two properties of the model make that merge unsound rather than merely inefficient. First, the derived profile is shared. PCOC assigns one amino acid profile to every branch declared convergent and a second profile to the remainder, then asks whether the declared branches all shifted to that same derived profile; the question is whether the lineages arrived at one destination, not whether each moved. Gains and reversals move toward opposite states, so no single derived profile describes both. At a site where the derived diurnal state favours one residue and the ancestral nocturnal state another, a merged run must either take the diurnal profile, which the reversal branches then contradict, or take the ancestral profile, in which case the convergent model collapses into the null. The cancellation is not a loss of sensitivity in weak data: the sharper the true signal in each direction, the more exactly the two offset. Second, a merged class would span 74 of the 118 branches on the species tree, 62.7 percent, leaving the background profile to be estimated from a minority of the tree and inverting the partition the model assumes.

Analysed separately, both directions are well powered at the calibrated threshold (Table 2), so keeping them apart costs no sensitivity.

**Table 2.**
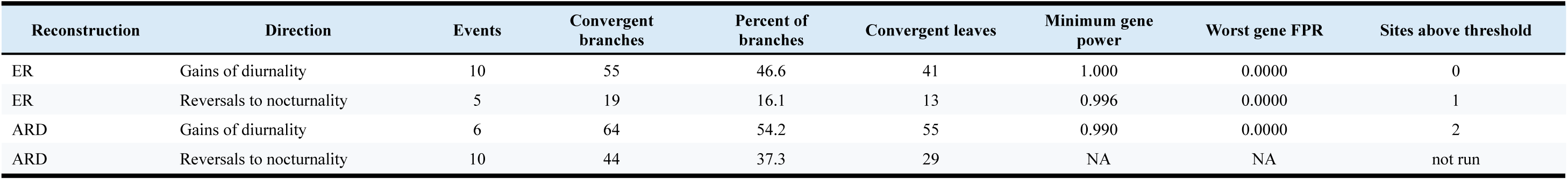
Convergent scenarios, calibrated detection power and outcome. Branch and leaf counts are on the 60-taxon species tree (118 branches). Power and false positive rate are the worst value across the 18 genes at the calibrated posterior threshold of 0.99.All sites above threshold were subsequently evaluated against alignment quality and reconstruction sensitivity. The ER-reversal site and one ARD-gain site were attributable to the ragged N-terminal alignment boundary, whereas the second ARD-gain site was specific to the less-supported ARD reconstruction and was not retained as a credible candidate. ER, equal rates; ARD, all rates different; FPR, false positive rate; NA, not available (scenario set not run).

For each transition, the convergent event comprises the transition branch plus every descendant branch remaining in the derived state. Descent terminates at any node that reverts, so nested reversals are excluded from the convergent group. Node identity was keyed by descendant tip set rather than node index, since node numbering differs between ape (Paradis and Schliep, 2019), ete3 (Huerta-Cepas et al. 2016)/IQ-TREE and PCOC and is not preserved under pruning. A node state table keyed by tip set was published once and read by every downstream consumer.

### Hemiplasy control

Gene and site concordance factors (Minh et al. 2020) were computed in IQ-TREE v3.1.1 in two separate runs against the fixed species tree. Gene concordance factors were computed from the 18 unconstrained maximum-likelihood per-gene trees (--gcf), so each branch’s gCF is the percentage of the 18 gene trees that recover the corresponding bipartition. Likelihood-based site concordance factors were computed from the partitioned locus alignments (-p, --scfl 100), drawing 100 quartet replicates per branch. The two per-branch tables were merged by IQ-TREE branch identifier, and each diel transition branch was matched to its concordance values by descendant tip set rather than by branch index.

Flagging followed a fixed conjunctive decision rule applied uniformly to all 15 transitions: a branch was flagged as hemiplasy-exposed only when gCF and sCFL both fell below 50 percent. Either criterion alone is commonly gene-tree estimation noise rather than genuine conflict, and a missing value on either axis never flagged. Concordance factors are undefined on tip branches, because a tip appears in every gene tree and there is no bipartition to conflict over, so the rule is evaluated only on the internal transitions.

### Convergence detection

PCOC (Rey et al. 2018) was run via pcoc_det.py from the carinerey/pcoc Docker image, taking each gene’s re-rooted fixed-topology tree, its trimmed protein alignment and the relevant convergent scenario. PCOC is distributed without numbered releases, so the image digest is the version of record; the build used here is: carinerey/pcoc@sha256:11ea18fb9b96e6bfa72bc694132745817ee77ec6b057fc8a2575698917a8598c. Gamma-distributed rate heterogeneity was enabled and the posterior reporting threshold was set to zero so that the full posterior distribution over sites was retained, with filtering applied afterwards rather than inside the detection step.

A site was called convergent only when its PCOC posterior reached 0.99. This cutoff was not optimised. Per-gene calibration (below) returned power 1.000 and a false positive rate of 0.0000 for every gene at every cutoff tested from 0.70 to 0.99, so the calibration does not identify a threshold; the strictest value in the saturated range was therefore adopted on the grounds that it costs no power. The same 0.99 cutoff governs every site-level decision reported here, including the comparison of a site’s posterior between reconstructions.

The posterior threshold was calibrated per gene on each gene’s own tree and scenario using pcoc_sim.py, with 100 simulated convergent sites and 100 null sites per profile couple and 10 profile couples per gene. The number of simulated events was set explicitly to each gene’s real event count, since the simulator otherwise samples a random subset of the supplied events. Calibration was performed independently for each of the three scenario sets analysed (Table 2; Supplementary Figure S1).

Two properties of the calibration bound its interpretation. First, pcoc_sim.py constructs the convergent shift with a OneChange model that conditions on a substitution occurring on every transition branch, so simulated power is conditional on a substitution having occurred rather than reflecting whether there was evolutionary time for one. Second, the detection step declares all events convergent regardless of the truth. To quantify the cost of that assumption, a partial-convergence sweep was run on one gene, CLOCK, chosen for having the full complement of taxa. For each k from 2 to 10, k of the 10 gain lineages were drawn and convergence was simulated in those lineages only, using pcoc_sim.py on the gene’s own tree with 100 convergent sites per profile couple and 2 profile couples, giving 200 planted sites per replicate. Detection then used pcoc_det.py with the full declared 10-event scenario, exactly as in the real analysis, so that the lineages in which nothing was planted remain declared convergent. Power at each k is the proportion of the planted sites whose posterior reaches the 0.99 cutoff.

Three independent draws of which lineages converge were taken at each k from 2 to 9. At k = 10 there is only one possible subset, so replicates would be identical draws and a single run was performed. Subsets were drawn from a seed fixed per k, so the sweep is reproducible. The sweep rests on a single gene and three replicates, and the replicate spread is wide at intermediate k, so it is reported as a characterisation of the design rather than as a precise power estimate.

### Site-specific fitness shifts

TDG09 v1.1.2 (Tamuri et al. 2009) was used to test, per site, whether amino acid fitness differs between nocturnal and diurnal lineages. Tree nodes were labelled from the published node state table rather than by propagating states from an assumed root, and the group argument order was set so that the nocturnal group is ancestral. Results were parsed from the FullResults block; per-site p-values were corrected within gene by the Benjamini-Hochberg procedure.

### Model-free residue screen

For each alignment column the statistic was the maximum over residues of freq(residue | diurnal) minus freq(residue | nocturnal), which equals 1.0 for a perfectly phenotype-diagnostic column. Columns with fewer than five ungapped residues in either group were skipped. Because diurnality is phylogenetically clumped, the null was generated not by shuffling tip labels but by re-placing clades of the observed sizes at random positions on the same gene tree, 1000 permutations per gene, so that the null retains the same autocorrelation structure and only the phenotype assignment is randomised.

This is a size-matched block permutation, a direct variant of the permulation principle of Saputra et al. (2021), which generates null phenotypes preserving both the number of foreground species and their phylogenetic relationships. Published permulations obtain that structure by simulating the trait on the tree under a fitted transition model and then binarising; the screen instead reproduces the observed clade size profile directly, because introducing a fitted evolutionary model would reimport the assumption the screen exists to avoid. The candidate pool constrains large blocks: in CLOCK, for instance, the tree offers 5 clades of size five against 59 of size one, so a permutation occasionally redraws an observed clade. That inflates the null toward the observed value and makes the test conservative, which is the safe direction for a null result and would need reporting were the result positive.

### Direct counting of convergent substitutions

Fitch parsimony (Fitch, 1971) was used to assign amino acids to internal nodes, so that a substitution on a branch is simply a difference between parent and child states. Because transition branches carry more substitutions of any kind than randomly chosen branches of matched clade size, raw counts are confounded. This is the confound identified by Thomas and Hahn (2015), whose reanalysis of echolocating mammals showed a reported excess of convergence to disappear once a null accounting for non-adaptive convergence was applied. Their null uses the correlation between convergent and divergent substitutions across species pairs; the null used here is a different implementation of the same principle. The test statistic was the proportion of sites changing in two or more independent lineages that changed to the same residue, and null branches were drawn matched both on clade size and on branch length to within a factor of two. Conditioning on opportunity addresses the symptom and matching branch length addresses the cause, so the two are reported together and are expected to agree.

Fitch parsimony treats the twenty amino acids as unordered and equally weighted, so a change from valine to isoleucine and one from valine to tryptophan both count as a single step. That is biologically false in two ways: the first requires one nucleotide substitution and the second at least two, and the first is chemically conservative while the second is not. The assumption is retained deliberately. Weighting the steps would require a substitution matrix, that is, a model of amino acid exchangeability, and this test exists as a model-free cross-check on the profile-based PCOC analysis; adopting a model of the same kind would remove its independence. The bias it introduces is also carried by the null, since observed and permuted branches are scored by the same counting, so it inflates the absolute number of apparent convergences without affecting the contrast on which the test rests.

The one place the matched null does not absorb it is the per-site test, where the null is matched on branches but not on how easily a residue can be reached. This is visible in the results below: half of the sites reaching q ≤ 0.05 converge on serine, which has six codons and sits one nucleotide from a large fraction of the other amino acids, so it is the residue most readily arrived at by chance.

Parsimony ties were resolved ten times per site with random tie-breaking to propagate ancestral state uncertainty. This is not standard practice; most applications commit to a single reconstruction. It was included because the choice is consequential: an earlier run of these data taking one arbitrary resolution per site returned 213 same-residue sites where the ten-resolution procedure reported below returns 201, a 6 percent shift arising entirely from which of several equally parsimonious histories happens to be selected. Each site received a permutation p-value from 2000 permutations and a Benjamini-Hochberg q-value across all genes.

Sites were additionally required to be phenotype-specific: the frequency of the convergent residue among diurnal species minus its frequency among nocturnal species was required to exceed 0.25. The threshold was set by judgment rather than calibrated, so two things about it are worth stating. It was fixed before the positive control existed and never subsequently adjusted, and the control, run two weeks later, provides independent evidence that it does not remove real signal: with the criterion applied, all 36 planted sites at ten converging lineages and all 18 at seven were still recovered, while false positives fell from 195 to 26. The cost falls on weak convergence, where recovery at three converging lineages drops from 35 of 36 to 23 of 36. Sensitivity of the result to the threshold is reported in Results. Without this criterion the test identifies homoplasy unrelated to diel activity, because a residue arising repeatedly across the tree will by chance fall on some transition branches, and the real transitions form a phylogenetically clustered set while the null scatters branches more widely.

This second criterion is an addition to the standard procedure, and it was introduced in response to a diagnosed failure rather than imposed in advance. Applying the permutation test alone returned 12 sites at q ≤ 0.05 whose residues were not diagnostic of the phenotype: the median frequency gap among them was 0.067, three were negative, meaning the residue was commoner in nocturnal than in diurnal species, and half were serine, the residue most easily reached by chance under the unordered counting described above. The clearest case, CLOCK site 675, is a column otherwise fixed for methionine in 54 of 59 species: valine appears in four, two diurnal and two nocturnal, and the two nocturnal carriers are marsupials, the lineages most distant from the placental diurnal transitions. A rare variant arising independently in four scattered lineages is homoplasy, not convergence toward a phenotype. The requirement that a convergent residue also be phenotype-diagnostic is implicit in the logic of Forward Genomics (Hiller et al. 2012), which likewise conditions a genomic signal on matching the phenotype pattern, but the frequency-gap form used here is specific to this study.

### Relative evolutionary rates

RERconverge v0.3.0 (Kowalczyk et al. 2019) was used to test gene-level association between relative evolutionary rate and diel activity, consuming the uniform-model fixed-topology trees. Relative rates were computed with a square-root transform. The foreground was defined as diurnal tip branches only, the conservative choice. Association was tested with a binary phenotype correlation requiring at least 10 species and 2 foreground branches per gene, and p-values were corrected across the 18 genes.

Because parametric p-values from phylogenetic association tests are miscalibrated under phylogenetic non-independence, the association was also assessed against a permulation null (Saputra et al. 2021), which generates null phenotypes preserving both the number of foreground species and their phylogenetic relationships. One thousand permulations were generated in species-subset-match mode, which regenerates the null separately against each gene’s own taxon set, and repeated in complete-case mode as a check. The foreground was constructed with terminal branches only in the permulations as in the observed test, so that the null corresponds to the analysis actually performed.

### Selection analyses

Codon-level analyses were run in HyPhy v2.5.93 (Kosakovsky Pond et al. 2020) on the codon alignments. Branch lengths were taken from the per-gene substitution trees rather than the time-calibrated species tree, and all-gap codon columns were removed, with a per-gene coordinate map retained so that post-filtering codon indices map back to protein and human residue coordinates. Contrast-FEL (Kosakovsky Pond et al. 2021) tested, per site, whether the ratio of nonsynonymous to synonymous substitution rates differs between the diurnal-transition branch set and the remainder of the tree, with correction applied both within gene and across genes. RELAX (Wertheim et al. 2015) was run to test for relaxed or intensified selection on the same branch set.

### Multi-method consensus

Per-site results from PCOC, TDG09 and Contrast-FEL were mapped through the trimmed to untrimmed to codon to human residue coordinate chain and integrated, with support from at least two methods required for a site to be considered high confidence.

### Positive control

Because a null result is uninformative unless the analysis is shown capable of detecting a signal, and because simulation under a method’s own model cannot detect an incorrectly constructed scenario, known convergent sites were planted into the real alignments and the entire analysis was repeated unchanged over the modified data, using identical trees, scenario files and scripts.

Sites were created by writing a residue absent at that column into the diurnal species descending from a chosen set of gain events. Candidate columns were required to be variable but not saturated (two to six residues present, at most 10 percent gaps). Ninety sites were planted across all 18 genes at three difficulty levels: all 10 gain events converging, 7 of 10, and 3 of 10. Each site was verified before retention: the planted residue had to be essentially absent from nocturnal species, the diurnal minus nocturnal frequency gap had to exceed 0.10, and at least two independent events had to contribute. Recovery was scored against a withheld key.

### Software and data availability

IQ-TREE v3.1.1; trimAl v1.5.rev1; corHMM v2.8; phytools v2.5.2; ape v5.8.1; RERconverge v0.3.0; PCOC (carinerey/pcoc container); TDG09 v1.1.2; HyPhy v2.5.93; pal2nal; miniprot; R v4.4.1; Python v3.11.3 with ete3 v3.1.3 and NumPy v1.25.2 (Harris et al. 2020). Figures were produced with Plotly (Plotly Technologies Inc. 2015) and exported as vector PDF. Analysis code, scenario files and result tables are available at the project repository: https://github.com/alikoraykoc/circadian_phylogeny

## Results

### Evolutionary scenario

Diel activity shows significant phylogenetic signal (Pagel’s λ = 0.671, p = 0.010). The equal-rates reconstruction recovers a nocturnal ancestor for the placental crown without that state being imposed, but weakly: the marginal posterior at the placental crown under ER is 0.604 nocturnal. Six marsupials are included as an outgroup, so the root of the sampled tree is the therian node, not the placental ancestor, and the posterior at the therian root under ER is 0.546 nocturnal. Both values are close to even, and the balanced 30:30 design pulls the reconstruction towards diurnality relative to real mammalian frequencies; the nocturnal state survives that pull but by a margin too narrow to treat as support for the nocturnal bottleneck hypothesis. The same reconstruction infers 10 independent gains of diurnality and 5 reversals to nocturnality across 15 transition branches, 10 terminal and 5 internal (Figure 1, Table 2).

**Figure 1.**
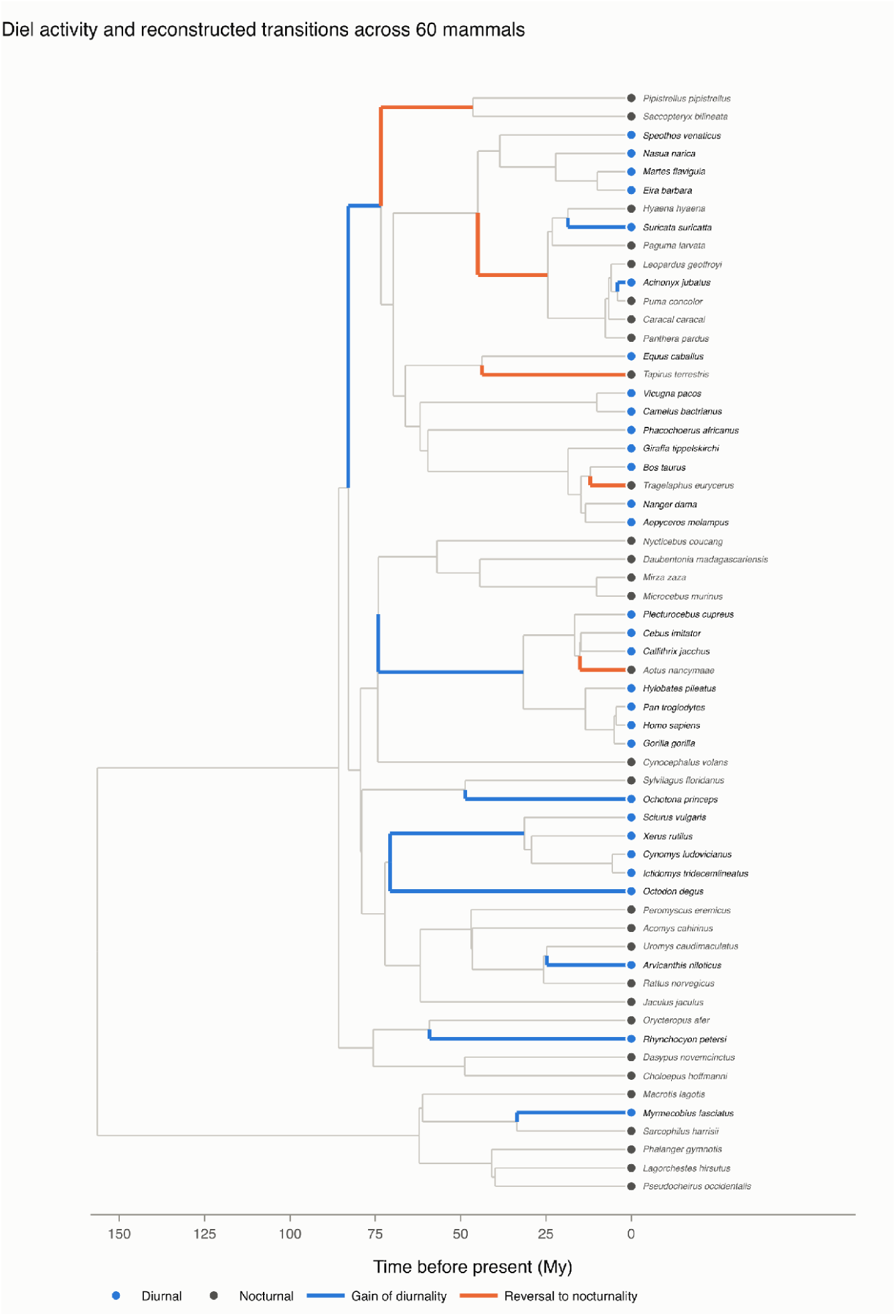
Diel activity and reconstructed transitions across 60 mammals. The fixed, time-calibrated mammalian supertree used for scenario construction is shown with observed diel states at the tips and transition branches inferred under the primary equal-rates ancestral-state model. The reconstruction identified 10 independent gains of diurnality and 5 reversals to nocturnality. Branch lengths are shown in millions of years.

The all-rates-different reconstruction instead places a diurnal ancestor at the placental crown, rooting diurnality at that 54-tip clade, and reframes the history as 6 gains and 10 reversals. Its margin is as narrow as the one it overturns: the placental crown under ARD is 0.468 nocturnal, while the therian root under ARD is 0.579 nocturnal. The two models therefore disagree only at the placental crown and agree at the sampled-tree root, and neither resolves the placental crown. This is the expected consequence of the deliberately balanced diel sample and is the reason the equal-rates model is treated as primary.

Ten of the 15 transitions sit on tip branches, where gene tree discordance cannot arise: a tip is present in every gene tree, so there is no bipartition to conflict over. Of the five internal transitions, none is weak on both concordance axes (gCF below 50 percent and sCFL below 50 percent) and none was flagged; three are weak on one axis (Table 4). The one worth naming is the largest gain of diurnality in the scenario, the branch subtending 24 taxa, where gCF is 38.9, the lowest of any transition, against sCFL of 75.7. Low gene concordance with high site concordance is the signature of gene tree estimation error rather than genuine conflict: individual genes carry too few informative sites to recover the clade, while the sites themselves support it strongly. That combination is precisely what the conjunctive criterion is designed to distinguish: flagging on gCF alone would have discarded the study’s largest convergent event because individual gene trees were poorly resolved despite strong site-level support. Gene tree discordance therefore cannot plausibly generate false convergence at any transition in this dataset.

### The analysis detects planted convergence

Ninety convergent sites planted into the real alignments were recovered as follows (Table 3, Figure 2).

**Figure 2.**
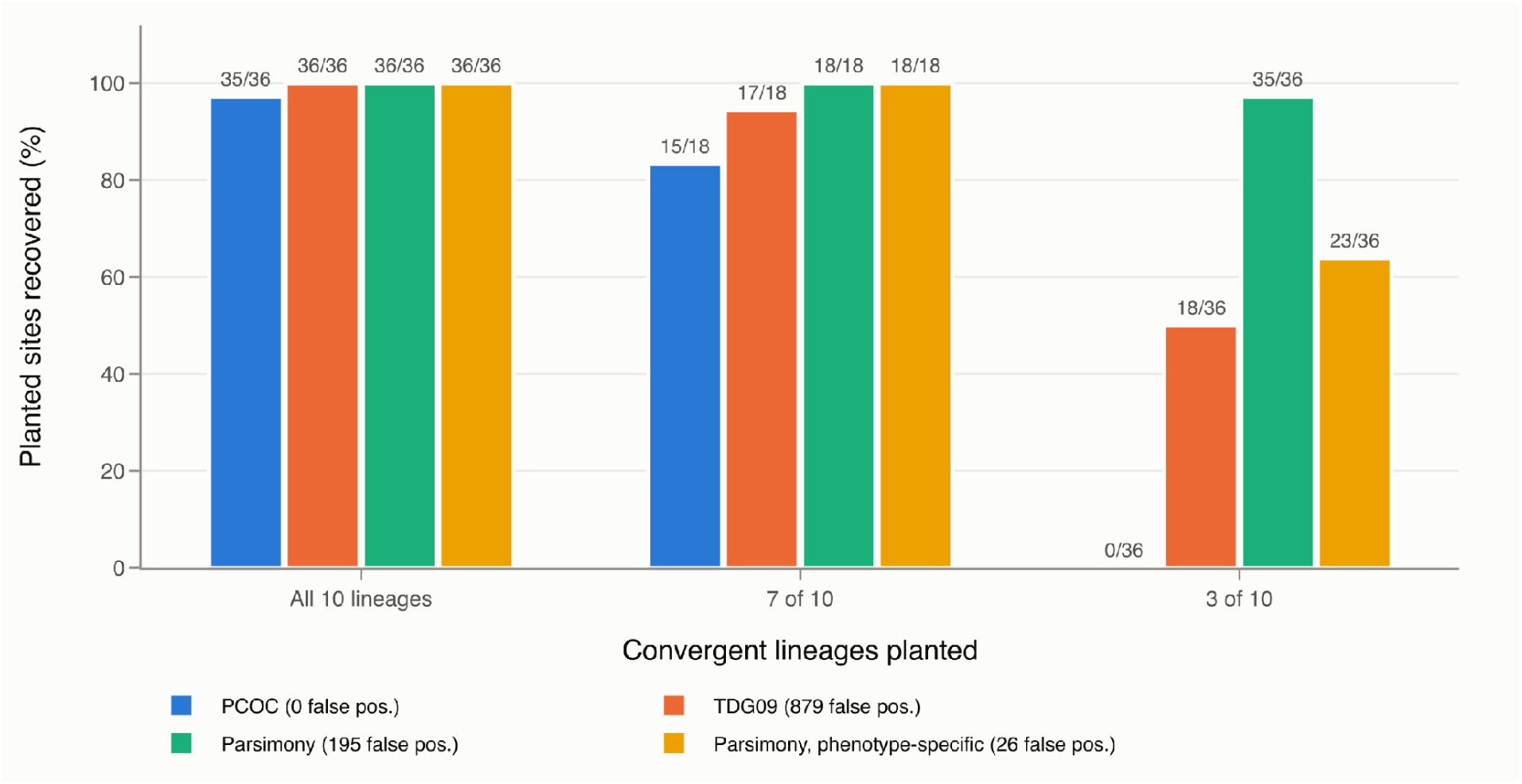
Recovery of planted convergent sites in the positive-control analysis. Ninety sites were planted into the empirical alignments at three levels of convergence: all 10 diurnality-gain lineages (n = 36), 7 of 10 lineages (n = 18), and 3 of 10 lineages (n = 36). Bars show the percentage recovered by PCOC, TDG09, the unfiltered parsimony test, and the phenotype-specific parsimony test; labels above the bars give the corresponding recovered-site counts. False-positive counts among unplanted sites are reported in the legend. PCOC calls required a posterior of at least 0.99; statistical site calls required q ≤ 0.05, and the phenotype-specific parsimony test additionally required a diurnal-minus-nocturnal residue-frequency difference greater than 0.25.

**Table 3.**
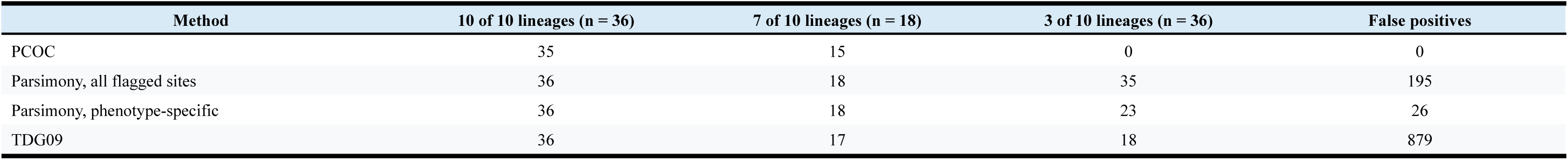
Positive control recovery by method and difficulty level, with false positive counts. Ninety convergent sites were planted into the real alignments at three difficulty levels, defined by how many of the 10 diurnal lineages converge at the site. Values are the numbers of planted sites recovered. False positives are unplanted sites flagged on the modified alignments; for TDG09 this corresponds to 22.8 percent of 3,852 testable sites. Parsimony results are shown both for all flagged sites and after requiring phenotype specificity.

| Method | 10 of 10 lineages (n = 36) | 7 of 10 lineages (n = 18) | 3 of 10 lineages (n = 36) | False positives |
| --- | --- | --- | --- | --- |
| PCOC | 35 | 15 | 0 | 0 |
| Parsimony, all flagged sites | 36 | 18 | 35 | 195 |
| Parsimony, phenotype-specific | 36 | 18 | 23 | 26 |
| TDG09 | 36 | 17 | 18 | 879 |

**Table 4.** Gene and site concordance at every internal transition branch. The remaining 10 of the 15 transitions are on tip branches, where a concordance factor is undefined because a tip appears in every gene tree. A branch is flagged as a hemiplasy risk only when gCF and sCFL are both below 50 percent, since either alone is usually gene-tree estimation error rather than genuine conflict. No transition is flagged. gCF, gene concordance factor; sCFL, site concordance factor (likelihood-based).

| Event | Direction | Descendant taxa | gCF | sCFL | Decisive gene trees | Weak axis | Flagged |
| --- | --- | --- | --- | --- | --- | --- | --- |
| 1 | gain | 24 | 38.9 | 75.7 | 18 | gCF | No |
| 3 | reversal | 8 | 50.0 | 42.0 | 18 | sCFL | No |
| 8 | gain | 8 | 55.6 | 60.8 | 18 | None | No |
| 2 | reversal | 2 | 56.2 | 48.1 | 16 | sCFL | No |
| 11 | gain | 4 | 58.8 | 55.4 | 17 | None | No |

PCOC flagged exactly 50 of 11,727 sites on the modified data, and all 50 were planted: 35 of 36 sites at which all 10 lineages converged, and 15 of 18 at 7 of 10, with no false positives. Signal therefore survives scenario construction, tree handling and detection intact.

The phenotype-specific parsimony test recovered all 36 sites at 10 lineages, all 18 at 7 of 10, and 23 of 36 at only 3 of 10, with 26 false positives. Requiring phenotype specificity reduced false positives from 195 to 26, an 87 percent reduction, at no cost to recovery at the two higher levels.

PCOC recovered none of the 36 sites at 3 of 10 lineages, as its design predicts. Sensitivity to convergence confined to few lineages therefore rests on the parsimony test, which retains approximately 64 percent recovery at that level.

TDG09 recovered 36 of 36, 17 of 18 and 18 of 36 respectively, but produced 879 false positives among 3,852 testable sites, a rate of 22.8 percent.

### No convergent signal accompanies gains of diurnality

Across all 18 genes, 648 sites changed in two or more independent diurnal lineages, and 201 of those changed to the same residue, an observed rate of 0.310 against a branch-length-matched null of 0.337 (Figure 3). No gene shows a rate-conditioned p-value below 0.282.

**Figure 3.**
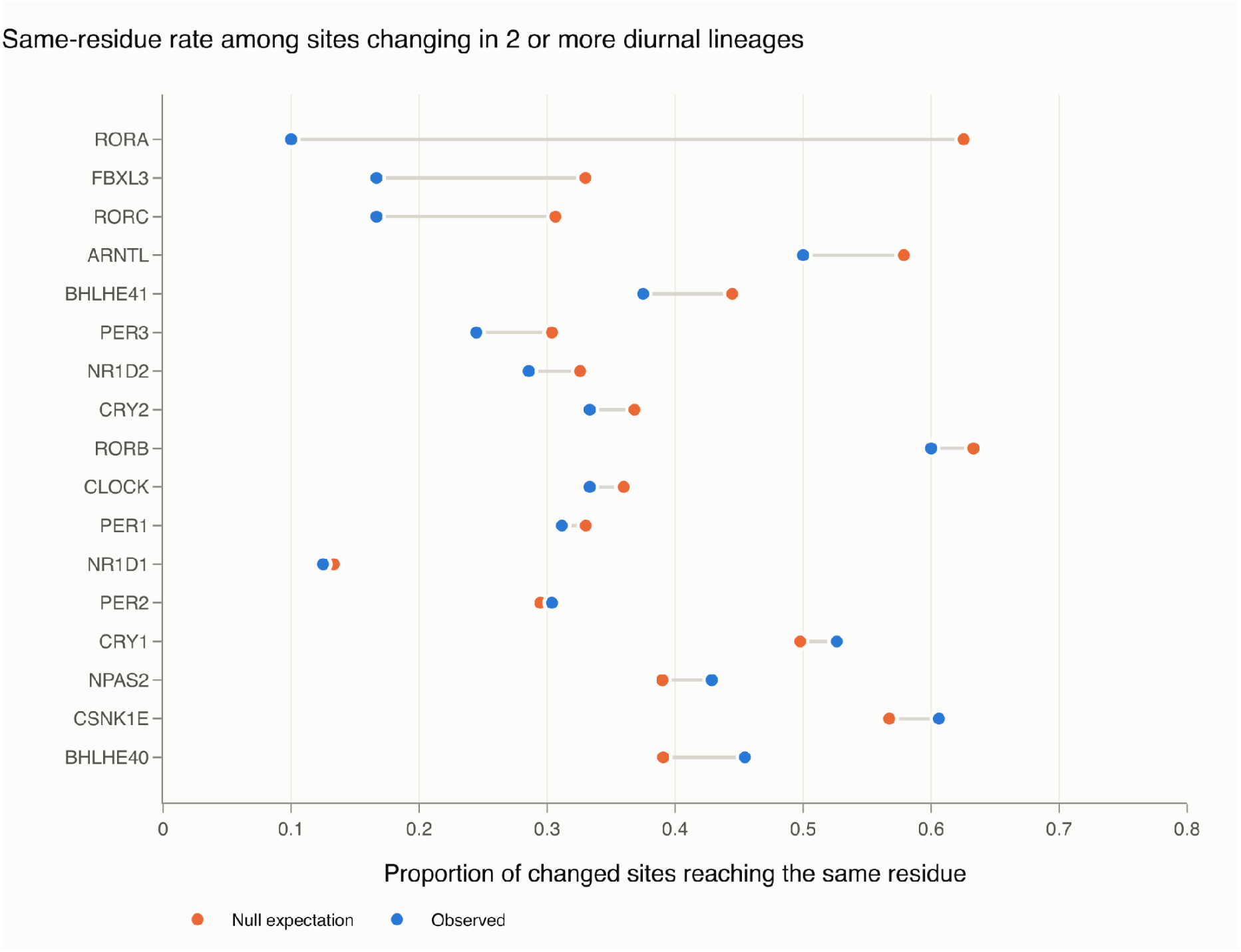
Observed same-residue convergence rates and matched null expectations by gene. For each gene, the observed value is the proportion of sites that changed independently in at least two diurnal lineages and reached the same amino acid. Null expectations were obtained from 2,000 permutations matching branches by descendant-clade size and branch length. CSNK1D is absent because it contained no site with substitutions in two or more independent diurnal lineages.

Twelve of the 201 same-residue sites reached q ≤ 0.05, but none is phenotype-specific. Nine of the twelve carry a residue as common in nocturnal as in diurnal species; the clearest example sits in a column otherwise fixed for methionine in 54 of 59 species and carries valine in two diurnal and two nocturnal species, a frequency gap of −0.002, with two of the four carriers being nocturnal marsupials. The largest diurnal minus nocturnal gap among the twelve is 0.232 and the median is 0.067, so no site is both statistically unusual and phenotype-specific.

The two criteria are close to orthogonal in these data. Of the 201 same-residue sites, 12 reach q ≤ 0.05 and 31 exceed the 0.25 phenotype-specificity gap, but no site belongs to both sets.

Because the 0.25 threshold was judgment-based, we report the result across a range of thresholds. The largest gap among the 12 statistically unusual sites is 0.232, so every threshold above that value returns zero, and the count rises only as the threshold is relaxed below it (Table 5).

**Table 5.**
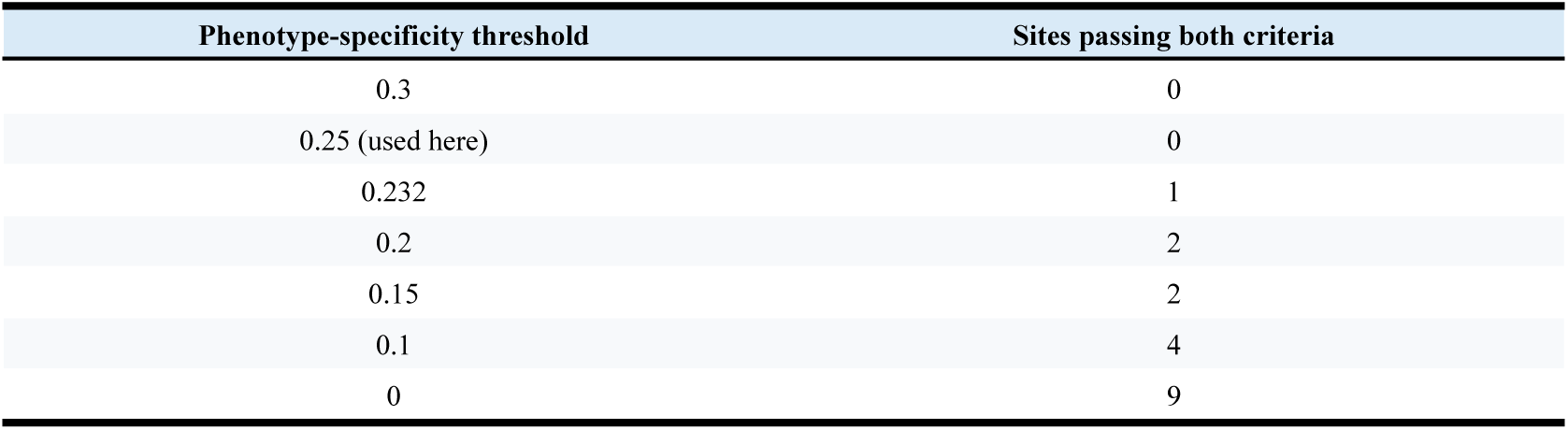
Convergent sites as a function of the phenotype-specificity threshold. Sites counted are those that are both statistically unusual (q ≤ 0.05) and phenotype-specific at the given threshold. The 0.25 cutoff was fixed by judgment before the positive control existed and never adjusted; the row between 0.25 and 0.20 is the largest gap observed among the statistically unusual sites, so every threshold above it returns zero.

At more permissive thresholds, only two sites survive: PER1 site 876 (serine, three events, gap 0.232) and BHLHE40 site 359 (serine, three events, gap 0.200). Both are weakly phenotype-associated by construction, since a gap of 0.2 means the residue is 20 percentage points commoner in diurnal than in nocturnal species, and both are serine. Three of the 12 have negative gaps and so fail at any positive threshold.

The claim this supports is therefore narrower than “no site passes” and more robust: across the plausible range of the threshold there is at most one weak candidate per gene in two genes, and none at the value chosen in advance.

The model-free residue screen agrees. Observed maximum diagnostic scores sit on the permutation null in all 18 genes and below it in several (RORB 0.223 against 0.305; CSNK1D 0.250 against 0.326). Only 2 of 11,727 columns exceed a diagnostic score of 0.6, against a comparable null expectation, and the lowest p-value (CRY1, 0.043) does not survive correction across 18 genes.

PCOC returned no site above threshold in any gene. The highest posterior observed across 11,727 columns is 0.098, and every other gene maximum is 0.003 or below. Calibration on these scenarios gives power 1.000 and a false positive rate of 0.0000 at every threshold tested, so this is not a marginal result.

### PCOC detects only near-universal convergence

Simulating convergence in only k of the 10 declared lineages and detecting with the full scenario shows power rising from 0.000 at k = 2 and 3, through 0.007 at k = 5 and 0.187 at k = 7, reaching 1.000 only at k = 10. These are powers at the calibrated posterior threshold of 0.99 used throughout; at the more permissive threshold of 0.90 the same points are 0.012 and 0.258. Supplementary Figure S2 shows both curves. They are identical for k = 2 to 4 and at k = 10, and differ most at k = 7, so the conclusion does not depend on which cutoff is read. The curve rests on one gene (CLOCK) with three replicates at each k from 2 to 9 and a single run at k = 10, where the subset of converging lineages is unique, Consequently, it characterises the method’s behaviour for CLOCK only and should not be interpreted as an across-gene power distribution. A direct check confirms the mechanism: at k = 2, detection under the declared 10-event scenario recovered 0 of 600 planted sites, while the same data detected under the correct 2-event scenario recovered 600 of 600.

The controlling variable is the number of falsely declared events rather than the quantity of sequence change available. Comparisons at matched declared-branch counts separate them: the two k = 2 replicates carrying 28 branches and 8 falsely declared events give power 0.000, whereas replicates of comparable size but fewer false events retain appreciable power, 0.100 at k = 6 with 25 branches and 4 false events, and 0.205 and 0.015 in the two k = 7 replicates with 26 branches and 3 false events. The replicate spread at k = 7 is wide, so this comparison establishes the direction of the effect and not its magnitude. PCOC’s model requires the derived profile on every declared branch, so each declared lineage that did not converge penalises the fit.

The simulation and the spike-in experiment disagree at the same value of k, and the disagreement bounds how the curve should be read. PCOC recovered 15 of 18 planted sites at 7 of 10 lineages, a recovery of 0.833 with a 95 percent Clopper-Pearson interval of 0.586 to 0.964, whereas the simulated curve gives 0.187 at k = 7. Both figures are taken at the same 0.99 posterior threshold, so the gap is not a threshold artefact, and it is not replicate noise either: the most favourable of the three replicates reaches only 0.340, below the lower bound of the spike-in interval. What differs is the strength of the planted signal. The simulation applies a probabilistic profile shift, whereas the planted sites carry an unambiguous novel residue in every diurnal descendant. PCOC’s sensitivity to partial convergence therefore depends on how clean the convergent substitution is, and the curve above describes performance for probabilistic profile shifts and does not establish a general power bound.

The PCOC result should therefore be read as excluding convergence shared by nearly all diurnal lineages, a narrower claim than excluding convergence generally. The general claim rests on the parsimony and residue-screen results, which declare no convergent set.

### Reversals to nocturnality

The reversal direction, tested here for the first time, returned 1 site of 11,727 above threshold (RORB, trimmed column 1, posterior 0.99999963); every other gene maximum is 0.119 or below and 11 of 18 are exactly zero.

That site is an alignment artifact. The trimmed alignment’s first column maps to untrimmed column 204, and 20 of the 60 sequences have their annotated protein begin at exactly that position, making the column an initiator residue for a third of the dataset and an internal residue for the rest. It carries 14 distinct residues, whereas the five columns immediately following each carry a single residue in 59 of 60 species. The 13 leaves descending from the five reversal events carry eight different residues, giving a best diagnostic score of 0.134 against a null expectation of 0.305 for that gene.

The reported posterior is carried by the one-change component, which is 0.9999994 at this site and is trivially satisfied at a hypervariable column. The profile-change component cannot be used to make this point. In the software build used here it returns exactly 0.500 at every site of every gene, because the likelihood of the no-one-change model is numerically invariant to which derived profile is declared, so the component equals its uniform prior by construction rather than by measurement (Supplementary Methods S1). The rejection of this site therefore rests on the alignment evidence above, which is independent of the posterior decomposition.

### Sensitivity to the ancestral-state model

Repeating the analysis under the all-rates-different reconstruction, for which calibration gives power 0.990 to 1.000 and a false positive rate of 0.0000, returned 2 sites of 11,727 above threshold, both in RORB and both in the same ragged N-terminal region (columns 1 and 2, posteriors 0.998 and 0.9996). Column 1 is the site discussed above and is rejected on the same alignment grounds, being hypervariable with 14 distinct states. Column 2 is not, and requires a different argument: it carries 8 distinct states and a best diagnostic score of 0.278, both inside the tolerances of the residue criteria. It is rejected because the signal is specific to this reconstruction. The posterior at the same site under the primary equal-rates reconstruction is 0.000380, four orders of magnitude below threshold, so the site is detected only under the ancestral-state model that fits worse by AIC and that infers a different ancestral activity pattern. Because the equal-rates model was designated primary on AIC in advance, this disposition does not depend on a choice made after seeing the result. All other gene maxima are 0.284 or below, and 12 of 18 are exactly zero.

This sensitivity analysis is weaker than the primary one: the ARD reconstruction declares 55 of 60 leaves and 54 percent of branches convergent, against 41 leaves and 47 percent under ER, leaving comparatively little ancestral contrast. The ARD result is consistent with the primary analysis, but it does not provide independent confirmation.

### Selection and rate association

Contrast-FEL identified no site of 15,349 with a significant difference in the nonsynonymous to synonymous rate ratio between the diurnal-transition branch set and the remainder of the tree, at a 5 percent false discovery rate either within gene or across genes; the smallest uncorrected p-value is 2.2 × 10⁻⁴. The within-gene correction is the operative one. With 15,349 tests the across-gene Benjamini-Hochberg procedure requires an uncorrected p-value below 3.3 × 10⁻⁶ at its smallest rank, which no site approaches, so every across-gene q-value is exactly 1.000 and that arm of the correction has no resolution at this dataset size.

RELAX did not run reliably on this dataset. Nine of 18 genes completed successfully, five failed after three attempts each during ancestral reconstruction on deeply divergent lineages, and the remaining four were not attempted. NPAS2 was the only nominally significant result among the nine completed genes (K = 0.352, q < 1 × 10⁻⁵), but it reproduced in only one of eleven runs, consistent with a flat likelihood surface. Because successful completion selected a non-random subset of genes, no inference is drawn from this analysis.

RERconverge identified no gene whose relative evolutionary rate is significantly associated with diel activity; all adjusted p-values exceed 0.67. The strongest raw signal is BHLHE40 (ρ = −0.178, p = 0.051, adjusted p = 0.677). The correlation sign is negative in 13 of 18 genes, indicating marginally slower relative rates in diurnal lineages, the opposite of the expectation under accelerated adaptive evolution, though far from significance.

The permulation null confirms this and shows the parametric p-values to have been anticonservative. Under 1000 species-subset-match permulations no gene reaches p = 0.05, the smallest permulation p-value being 0.114 for BHLHE40 against its parametric 0.051, and the smallest adjusted value 0.728. The parametric p-value is smaller than the permulation p-value in 10 of the 18 genes, in two cases substantially (NPAS2 0.127 against 0.673; ARNTL 0.137 against 0.361). Complete-case permulations give the same picture (smallest p 0.104, smallest adjusted 0.764). The one gene whose parametric value approached conventional significance therefore does not survive a null that accounts for phylogenetic structure. Calibration therefore provides additional support for the negative result.

### Cross-method integration

TDG09 flagged 885 of 3,820 testable sites at a 5 percent false discovery rate, 23.2 percent, a figure exceeding every other method by orders of magnitude. No flagged site is supported by any second method, and the high-confidence consensus set is empty (Figure 4). The positive control provides the direct comparison: on modified alignments the same procedure produced false positives at 22.8 percent (879 of 3,852 testable sites), statistically indistinguishable from its real-data flagging rate of 23.2 percent. Its recovery of planted sites is high, so the limitation is specificity rather than sensitivity.

**Figure 4.**
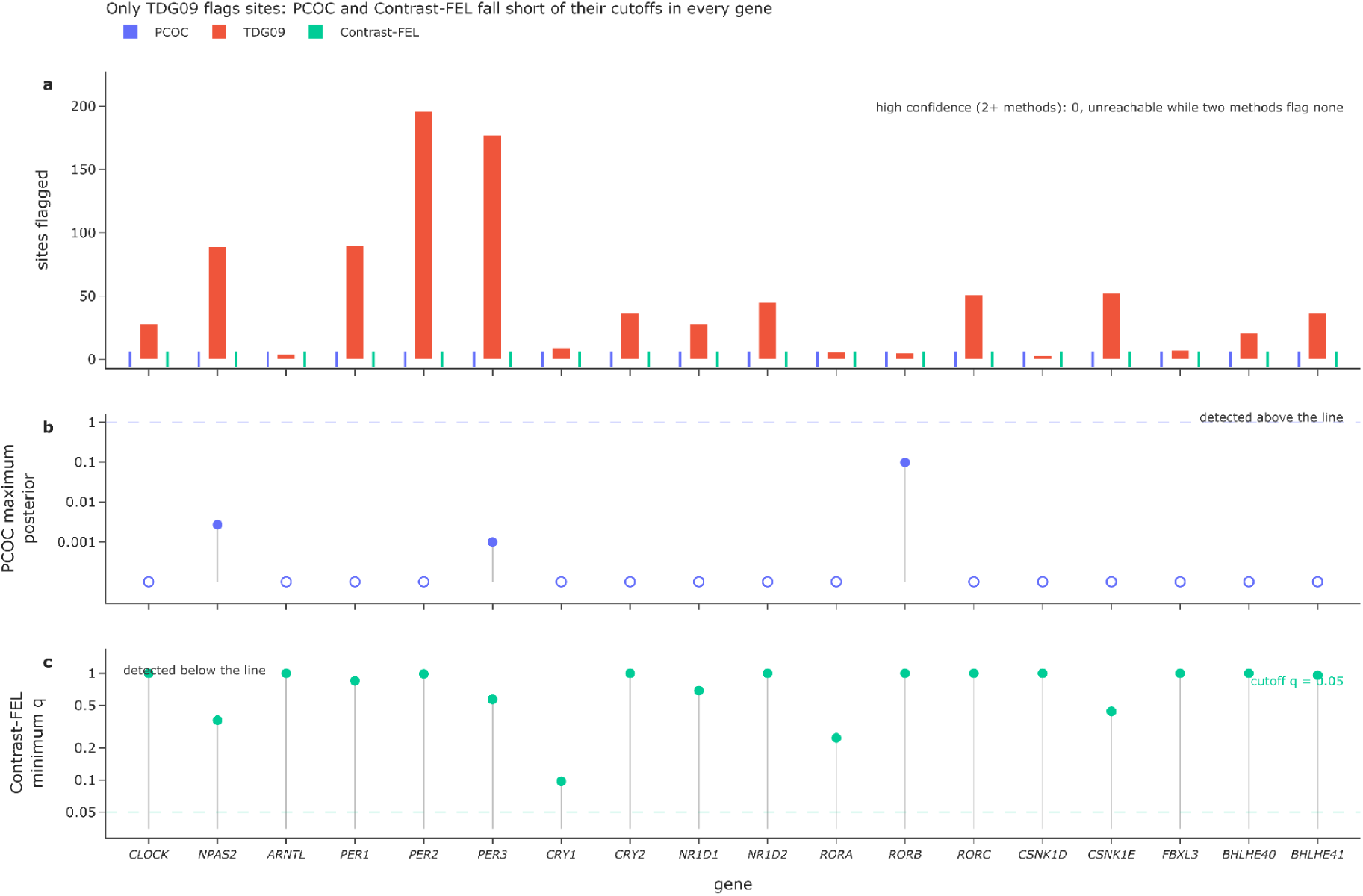
Per-gene site calls and distance from the PCOC and Contrast-FEL decision thresholds. (a) Final site calls per gene for PCOC, TDG09 and Contrast-FEL in the primary ER-gain analysis. TDG09 flagged 885 of 3,820 testable sites (23.2%), whereas PCOC and Contrast-FEL returned no calls. Methods returning zero calls are marked on the baseline. No TDG09 call received support from a second site-level method, and the high-confidence consensus set was therefore empty. (b) Maximum PCOC posterior per gene relative to the 0.99 cutoff, shown on a logarithmic scale. Across 11,727 scored alignment columns, the highest posterior was 0.0979 in RORB; maxima for 15 genes underflowed to zero and are shown at the plotting floor. (c) Minimum within-gene Contrast-FEL q value per gene relative to the 0.05 cutoff, shown on a logarithmic scale. Across 15,349 sites, the smallest q value was 0.0975 in CRY1.

### Genes with insufficient opportunity

Two genes cannot address the question. CSNK1D contains no position at which two or more independent diurnal lineages changed, and ARNTL contains two, against expectations of 2.0 and 2.1 usable sites respectively from branch lengths alone. CSNK1D also has the lowest taxon occupancy (40 of 60). Their null results are uninformative rather than negative and are reported separately throughout.

## Discussion

### No shared coding signature across repeated temporal-niche transitions

We found no consistent evidence that independent transitions between nocturnal and diurnal activity were accompanied by the same changes in the protein-coding sequences of the 18 clock-associated genes examined. PCOC detected no credible convergent site associated with gains of diurnality or reversals to nocturnality. Direct substitution counting likewise found no excess of same-residue convergence on diurnal-transition branches relative to the branch-length-matched null, and none of the statistically unusual substitutions also met the additional phenotype-specificity criterion. The model-free residue screen identified no gene-level association after correction, Contrast-FEL detected no significant site-wise shift in selective pressure, and RERconverge found no association between diel activity and relative evolutionary rate under the phylogenetically informed permulation null. The cross-method consensus set was therefore empty.

This agreement is informative because the methods were designed to detect different forms of molecular association. PCOC tests for a shared shift in amino acid preferences, direct counting identifies repeated substitutions to the same residue, the residue screen searches for phenotype-diagnostic sequence states without specifying an evolutionary model, Contrast-FEL tests for site-specific differences in nonsynonymous relative to synonymous substitution rates, and RERconverge operates at the level of gene-wide evolutionary rates. Their joint result argues against a single strong coding signature shared broadly across the sampled transitions. It does not imply that every transition followed the same molecular path without leaving a detectable trace. Phenotypic convergence can arise through different substitutions, different genes or different regulatory mechanisms in different lineages, even when the resulting phenotype is similar (Natarajan et al., 2016).

Not every analysis contributed equally to this conclusion. TDG09 classified 885 of 3,820 testable sites as significant, but none was supported by another method. Moreover, its false-positive rate in the planted-signal experiment was 22.8%, nearly identical to the 23.2% of empirical sites it flagged. Its empirical calls therefore cannot be distinguished from the method’s background error rate in these data. RELAX also failed to provide an interpretable result: only a subset of genes converged reliably, and the apparent signal for NPAS2 was not reproducible across repeated runs. No inference about relaxation or intensification of selection is consequently drawn from RELAX. These outcomes should be distinguished from negative results produced by analyses that ran reliably and were successfully calibrated.

### Detection limits of the analyses

The planted-signal experiment defines the strongest empirical basis for interpreting the negative result, but it does not provide a single measure of power for the entire study. Ninety sites were modified at three levels of convergence. PCOC recovered 35 of 36 sites modified across all 10 diurnality-gain events and 15 of 18 sites modified across 7 of the 10 events, with no false positives, but recovered none of the 36 sites modified across only 3 events. The phenotype-specific parsimony test recovered all sites at 10 and 7 events and 23 of 36 sites at 3 events, although with 26 false positives. The pipeline can therefore detect strong site-specific convergence shared across most transition lineages, while the parsimony analysis retains some sensitivity when convergence is restricted to a minority of them.

The additional PCOC simulations show that sensitivity depends not only on how many lineages converge but also on the form and strength of the signal. Cleanly planted novel residues were often recovered when present in 7 of 10 lineages, whereas probabilistic shifts in amino acid profiles were difficult to detect unless nearly all declared lineages shared them. The absence of a PCOC signal therefore weighs most strongly against clear and broadly shared convergence. It provides much weaker evidence against heterogeneous or partial convergence. More general power estimates would require replicated simulations spanning different genes, effect sizes, foreground proportions and substitution models (Rey et al., 2018).

The number of reconstructed phenotypic transitions provides useful context but is not itself a power estimate. The primary ancestral-state reconstruction identified 10 gains of diurnality and 5 reversals to nocturnality on particular branches, whereas Fitch parsimony assigned the complete tree a minimum score of 14 changes. These quantities answer different questions: the former describes one model-based reconstruction of transition branches, while the latter is the minimum number of changes compatible with the observed tip states. After gene-specific taxon pruning, minimum parsimony scores ranged from 9 to 14. These values describe how much transition replication remained in each gene, but statistical power also depends on topology, branch lengths, alignment length, missing-data structure and the distribution of the molecular signal.

### Sampling, phenotype assignment and gene coverage

Taxon sampling was jointly constrained by the availability of reliable behavioural evidence and genomic resources suitable for sequence recovery. The 60 species are therefore not a random sample of mammalian temporal diversity and disproportionately represent lineages with better genomic coverage. The initial 30:30 balance between diurnal and nocturnal species prevents either phenotype from dominating the analysis numerically, but it does not ensure equivalent phylogenetic representation. Deliberate balancing may also affect ancestral-state reconstruction when the sampled frequencies differ substantially from those in mammals as a whole.

The equal-rates reconstruction was selected as the primary scenario and recovered a nocturnal ancestor for the placental crown, at a marginal posterior of 0.604, so it is consistent with the broader evidence for a mammalian nocturnal bottleneck without providing independent support for it. The alternative all-rates-different model produced a markedly different history, including a diurnal ancestor at that node and a much larger declared convergent class; its margin is as narrow as the one it overturns, 0.468 nocturnal against 0.604, and the two models agree that the therian root is nocturnal (0.546 under ER, 0.579 under ARD), so the disagreement concerns the placental crown alone and is reported here rather than resolved. Repeating the analysis under this alternative scenario nevertheless produced no credible convergent site: the detected RORB positions were attributable either to a ragged alignment boundary or to dependence on the less-supported reconstruction. The main sequence-level conclusion was therefore not created by one particular ancestral-state reconstruction, although the weaker contrast under the alternative model limits the independence of this sensitivity analysis.

Diel activity is also more continuous and environmentally responsive than a binary classification implies. Several species show seasonal, geographical or population-level variation, and cathemeral or crepuscular activity cannot always be represented adequately as either diurnal or nocturnal. The confidence tiers and supporting behavioural evidence reported in Table S1 make this uncertainty visible, but they cannot remove classification error. Such error would generally reduce agreement among independently evolved lineages and could obscure a weak convergence signal.

Sequence loss introduced a related but distinct limitation. Per-gene diurnal-to-nocturnal balance ratios remained between 0.77 and 1.00 after quality control, but repeatedly missing taxa were mildly skewed towards the nocturnal class and clustered phylogenetically. Similar numbers of diurnal and nocturnal species therefore do not guarantee equivalent numbers of independent evolutionary transitions. This is why coverage and retained parsimony scores were evaluated separately for every gene.

The codon alignments were smallest for BHLHE40, CSNK1D, CSNK1E and PER3, which retained 38, 32, 39 and 36 species, respectively. CSNK1D requires particular caution: its protein alignment retained 40 species, its codon alignment retained 32, and its pruned topology had the lowest minimum parsimony score, 9 rather than 14. It also contained no site at which two or more independent diurnal lineages underwent a substitution. ARNTL contained only two such sites. For analyses requiring repeated substitutional opportunity, the null results for CSNK1D and ARNTL are therefore uninformative rather than evidence of absence.

Four taxa assigned to the diurnal class—Equus caballus, Bos taurus, Camelus bactrianus and Vicugna pacos—are domesticated, and their species-level activity assignments are uncertain (Table S1). Their observed behaviour may be influenced by feeding schedules, housing and artificial lighting rather than representing an evolved chronotype. Maor et al. (2017) classified E. caballus as cathemeral and did not include the other three species. Because these taxa were retained in the analyses, their classification remains a limitation of the present dataset. Any resulting misclassification would be expected to reduce correspondence between diel activity and sequence evolution, potentially obscuring a weak association.

The present sample was also too small to estimate the independent effects of several correlated life-history traits simultaneously. Body mass, longevity, hibernation and other ecological variables may influence protein evolutionary rates and may be correlated with diel activity. They could therefore mask or mimic weak rate associations. The failure to detect an activity-associated rate shift should not be treated as evidence that these covariates have no effect.

### Molecular changes outside recurrent amino acid convergence

The biological scope of the study is restricted to the coding sequences of a targeted panel of 18 clock components and direct regulators. The analyses do not directly test cis-regulatory variation in promoters and enhancers, tissue-specific expression, transcript or protein abundance, post-translational regulation, or changes elsewhere in the pathways connecting the oscillator to behaviour and physiology. Although the codon models use synonymous substitutions when estimating selective regimes, they were not designed to detect phenotype-associated changes in codon usage, mRNA secondary structure or translation efficiency. The study also does not cover every gene that has been described as part of, or closely connected to, the mammalian circadian system.

The distinction is particularly relevant in light of the experimental results of Beale et al. (2026). They found that cells from diurnal and nocturnal mammals entrained to temperature cycles in opposing phases and differed in the thermal sensitivity of protein synthesis, phosphorylation and circadian timing. Their comparative genomic analyses implicated broader signalling networks, including the mTOR and WNK pathways, and pharmacological inhibition of mTOR shifted nocturnal mouse cells, tissues and behaviour towards a more diurnal-like response. They also showed that the thermal response of PER2 translation was partly intrinsic to its coding region and was altered by codon optimisation, although PER2 was insufficient on its own to account for the nocturnal–diurnal switch.

These findings do not contradict the absence of recurrent amino acid convergence in our clock-gene panel. Instead, they identify mechanisms, polygenic rate changes across broader pathways, translational regulation, codon composition and differences in cellular signalling, that need not produce the same amino acid substitution at homologous sites in independent lineages. They also caution against describing chronotype evolution as occurring exclusively downstream of the oscillator: properties of PER2 itself may contribute, even if they do not take the form tested most directly here.

## Conclusion

Our results exclude a narrower class of explanations than an unqualified statement of “no molecular convergence” would imply. Under the taxon sampling, genes, evolutionary scenarios and decision thresholds used here, we found no strong and broadly shared coding-sequence signature that repeatedly accompanied mammalian transitions between nocturnal and diurnal activity. The result is supported by several complementary and calibrated analyses, but sensitivity declines for weak, lineage-restricted or gene-specific effects, particularly in genes with limited taxon coverage or substitutional opportunity.

The findings do not show that core clock proteins are evolutionarily irrelevant, that they cannot contribute to individual chronotype transitions, or that chronotype lacks a molecular basis. Rather, they indicate that repeated transitions between mammalian temporal niches were not generally achieved through one common set of detectable coding changes across these 18 genes. Disentangling the genetic architecture of mammalian chronotypes will likely require investigating cis-regulatory elements, post-transcriptional dynamics, and broader signalling cascades beyond the core oscillator.

## Data availability

Supplementary Table S1. contains the curated species-level diel-activity dataset, including the evidence, confidence, assignment qualifications and source references for each species. Supplementary Table S2 contains the species × gene provenance map for the 1,040 retained proteins, linking each record to either its annotated protein accession or its miniprot query and target contig. Additional data available in the project repository include the candidate-species pool and exclusion reasons, the reference topology, protein and codon alignments, verified unaligned coding sequences, per-sequence quality-control reports, per-gene class composition and balance ratios, and per-gene parsimony scores. Species-label equality and set membership were checked in both directions among the trait table, reference tree and each alignment. Analysis code, scenario files, processed result tables, and the data underlying all figures are available at the project repository: https://github.com/alikoraykoc/circadian_phylogeny.

## Author contributions

Alper Kaan Selçukoğlu led data collection, sequence retrieval, and data curation. Ali Koray Koç led the computational and statistical analyses. Both authors contributed equally to the interpretation of the results and to the writing, review, and revision of the manuscript. Both authors approved the final version and share first authorship.

## Declaration of Use of AI

During this study, the authors used Anthropic’s Claude and OpenAI’s ChatGPT as analytical and editorial assistants. These tools were used to identify potential errors in the analysis pipeline, support the cross-checking of chronotype assignments against the primary literature, and assist with language editing. All analyses were designed, implemented, interpreted, and verified by the authors. All literature citations, accession numbers, and numerical results reported in the manuscript were checked by the authors against the original sources and analysis outputs. The authors take full responsibility for the content of the manuscript.

## Supporting information

Table S2 Curated species-level diel-activity dataset

Table S1 Species x gene sequence-provenance map

