## Supplementary Information for "Testing for Shared Molecular-Evolutionary Signatures of Diurnality and Nocturnality in Mammalian Circadian Clock Genes"

##### Contents

Supplementary Methods S1  Examination of the PCOC posterior decomposition

Figure S1  PCOC power calibration on the empirical trees and scenarios

Figure S2  PCOC power under partial convergence for CLOCK

Table S1  Curated species-level diel-activity dataset  supplied separately as XLSX

Table S2  Species by gene sequence-provenance map  supplied separately as CSV

#### Supplementary Methods S1  Examination of the PCOC posterior decomposition

PCOC was run with pcoc_det.py from the carinerey/pcoc container (image digest sha256:11ea18fb9b96e6bfa72bc694132745817ee77ec6b057fc8a2575698917a8598c). The reporting threshold was set to zero so that the complete site-wise posterior output and its component values were retained. The diagnostic described here was performed on these retained outputs and did not alter the prespecified 0.99 calling threshold.

The profile-change component was examined across all analysed sites and genes. In the software build used here, it was exactly 0.500 at every site. Inspection of the corresponding likelihood terms showed that the likelihood of the model without the one-change constraint was numerically invariant to the declared derived profile. With equal prior odds, identical likelihoods return a posterior of 0.500. The profile-change component therefore remained at its uniform prior in these runs and supplied no site-specific evidence.

For the first trimmed RORB column discussed in the main text, the one-change component was 0.9999994. This component asks whether a substitution occurred on each declared transition branch and can consequently become large at a highly variable column. The column maps to untrimmed position 204, is the initiator residue in 20 of 60 sequences but an internal residue in the remainder, and contains 14 amino-acid states. The site was therefore rejected from biological interpretation on the independent alignment evidence, not on the uninformative profile-change component. This diagnostic did not change any site call or downstream consensus rule.

#### Figure S1

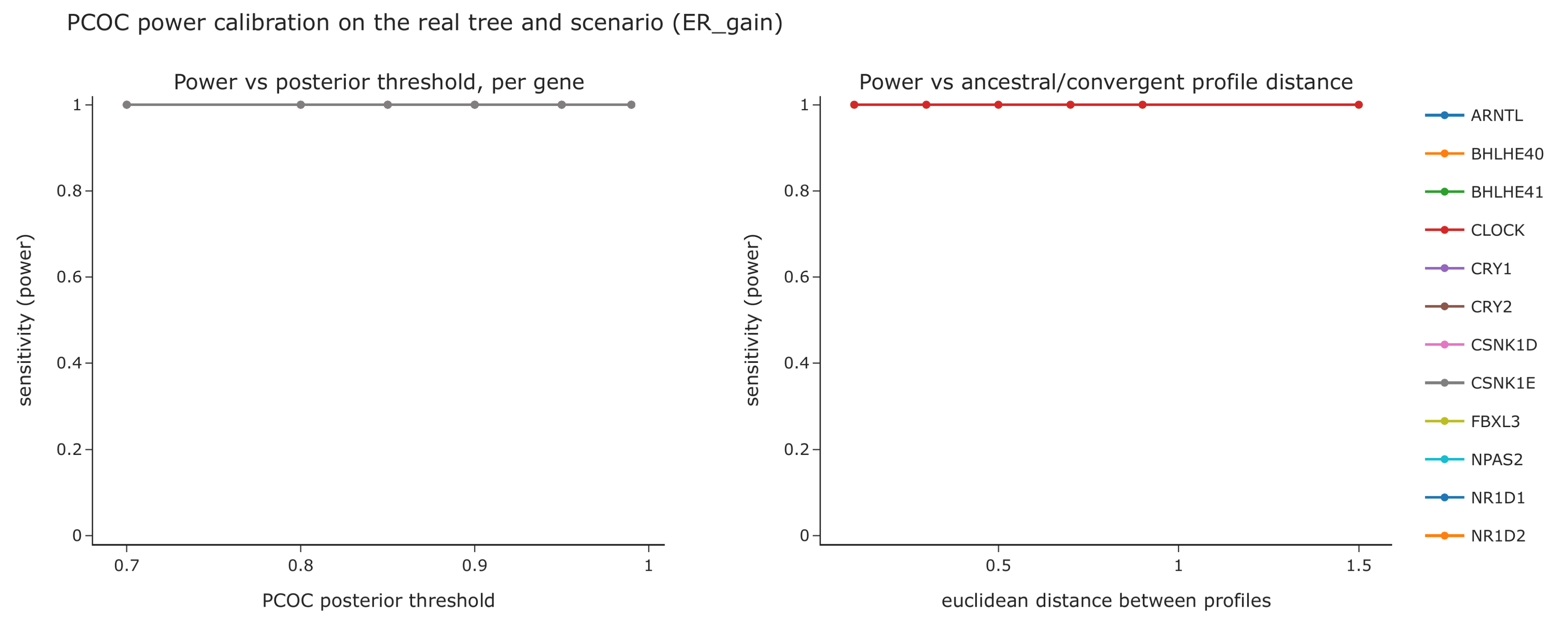


**Figure S1. PCOC power calibration on the empirical gene trees and convergence scenarios. The three pages show equal-rates** gains of diurnality (ER_gain), equal-rates reversals to nocturnality (ER_reversal), and all-rates-different gains of diurnality (ARD_gain). Left panels show per-gene sensitivity across posterior cutoffs from 0.70 to 0.99; right panels show sensitivity against the Euclidean distance between the simulated ancestral and convergent amino-acid profiles. Each calibration used the gene-specific tree and empirical scenario, 100 convergent and 100 null sites per profile pair, and 10 profile pairs per gene, with the simulated event count fixed to the empirical count. Under the primary ER reconstruction, power was 1.000 and the false-positive rate was 0.0000 across the tested cutoffs. Under ARD gains, power ranged from 0.990 to 1.000 and the false-positive rate was 0.0000.

**Figure S1 continued**

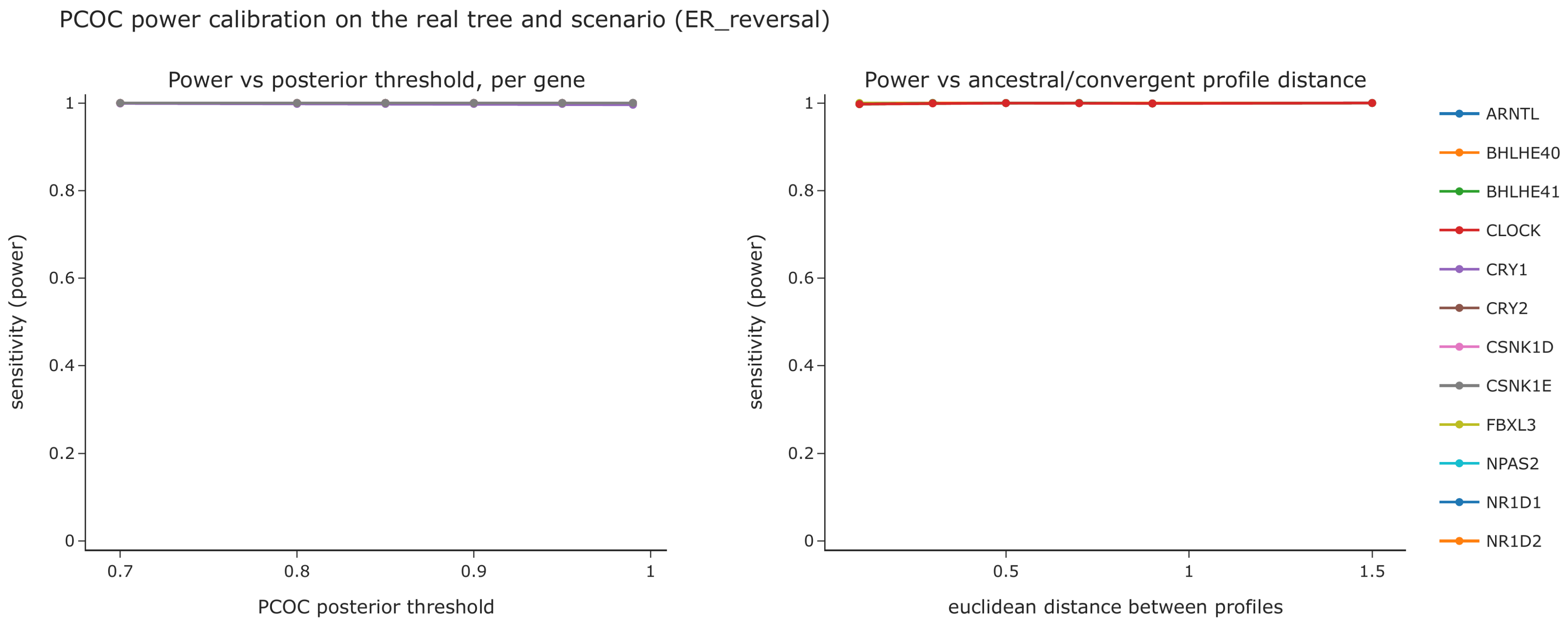

**Figure S1 continued**

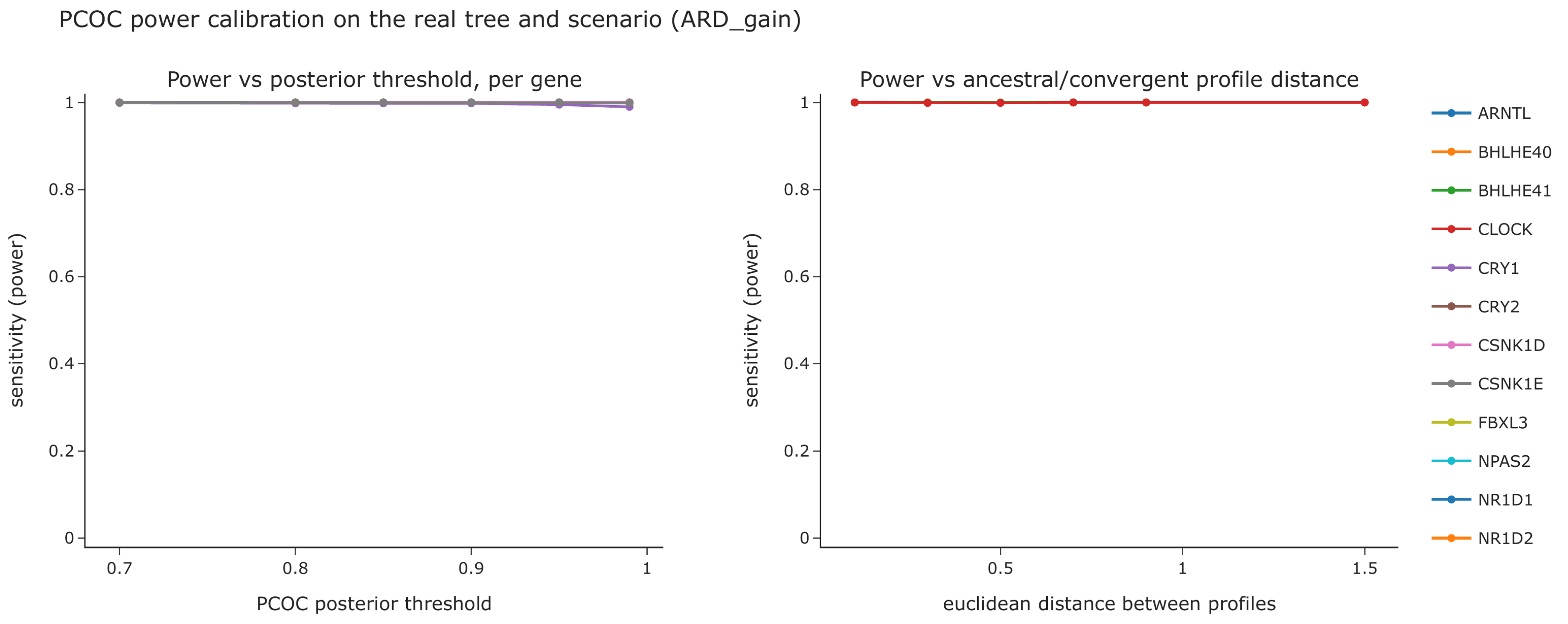

### Figure S2

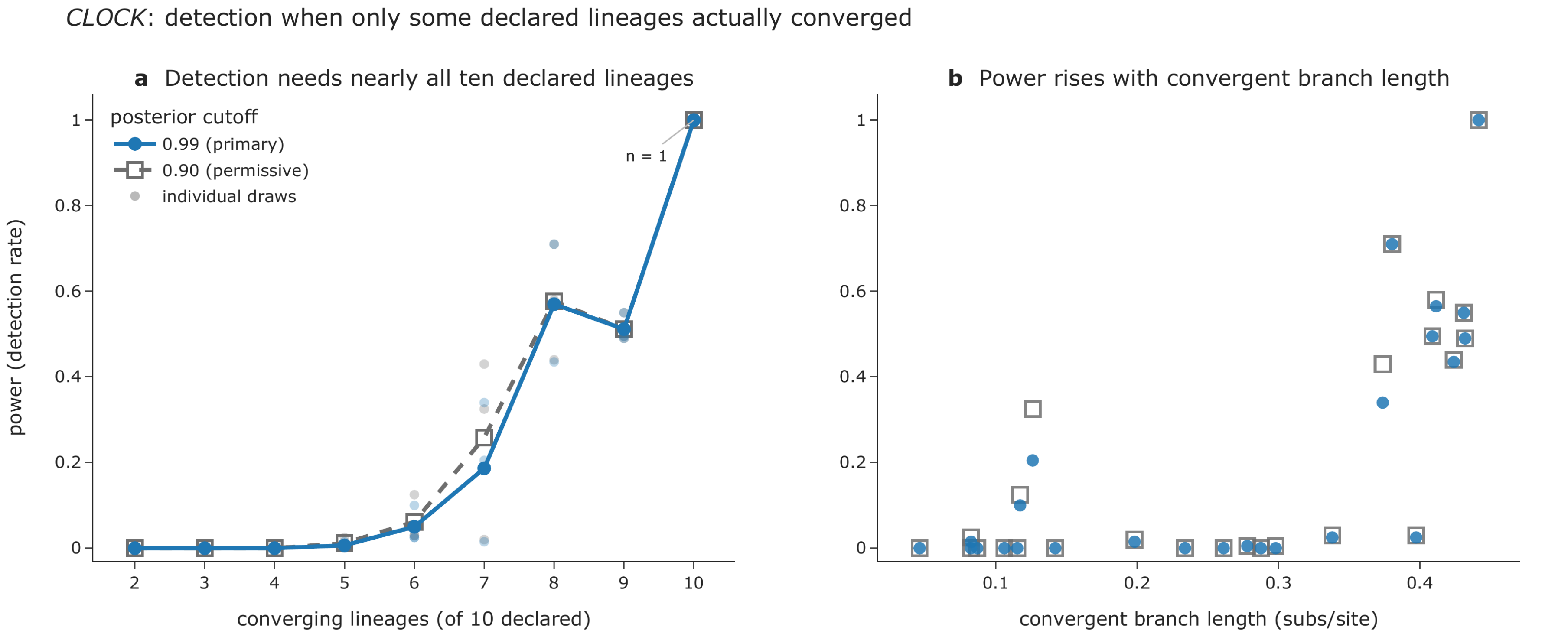


**Figure S2. PCOC power when only some declared lineages actually converged, for CLOCK. Detection always uses the full** declared 10-event scenario, as in the empirical analysis. Both posterior cutoffs are shown: 0.99, used throughout this study, and the permissive 0.90. (a) Power against the number of converging lineages; lines are means over three independent draws of which lineages converge at each k from 2 to 9, faint marks are the individual draws, and k = 10 is a single run because the converging set is then unique. (b) The same replicates against the total branch length of the converging lineages, which rises with k and is therefore not an independent predictor of power.
